# A novel inducible von Willebrand Factor Cre Recombinase mouse strain to study microvascular endothelial cell-specific biological processes *in vivo*

**DOI:** 10.1101/2023.07.24.550419

**Authors:** Dinesh Yadav, Jeremy A. Conner, Yimin Wang, Thomas L. Saunders, Eroboghene E. Ubogu

**Author notes:** Address all correspondence to: Dr. Eroboghene E. Ubogu Division of Neuromuscular Disease Department of Neurology University of Alabama at Birmingham 1720 7th Avenue South, Sparks Center Suite 200 Birmingham, AL 35294-0017 United States of America, Facsimile Number.

## Abstract

Mouse models are invaluable to understanding fundamental mechanisms in vascular biology during development, in health and different disease states. Several constitutive or inducible models that selectively knockout or knock in genes in vascular endothelial cells exist; however, functional and phenotypic differences exist between microvascular and macrovascular endothelial cells in different organs. In order to study microvascular endothelial cell-specific biological processes, we developed a Tamoxifen-inducible von Willebrand Factor (vWF) Cre recombinase mouse in the SJL background. The transgene consists of the human vWF promoter with the microvascular endothelial cell-selective 734 base pair sequence to drive Cre recombinase fused to a mutant estrogen ligand-binding domain [ERT2] that requires Tamoxifen for activity (CreERT2) followed by a polyadenylation (polyA) signal. We initially observed Tamoxifen-inducible restricted bone marrow megakaryocyte and sciatic nerve microvascular endothelial cell Cre recombinase expression in offspring of a mixed strain hemizygous C57BL/6- SJL founder mouse bred with mT/mG mice, with >90% bone marrow megakaryocyte expression efficiency. Founder mouse offspring were backcrossed to the SJL background by speed congenics, and intercrossed for >10 generations to develop hemizygous Tamoxifen-inducible vWF Cre recombinase (vWF-iCre/+) SJL mice with stable transgene insertion in chromosome 1. Microvascular endothelial cell-specific Cre recombinase expression occurred in the sciatic nerves, brains, spleens, kidneys and gastrocnemius muscles of adult vWF-iCre/+ SJL mice bred with Ai14 mice, with retained low level bone marrow and splenic megakaryocyte expression. This novel mouse strain would support hypothesis-driven mechanistic studies to decipher the role(s) of specific genes transcribed by microvascular endothelial cells during development, as well as in physiologic and pathophysiologic states in an organ- and time-dependent manner.

## Introduction

Vascular biologists have generated multiple mouse strains in different genetic backgrounds to tightly regulate gene deletion, or express transgenic genes in the murine endothelium over the past 30 years. These strains facilitate *in vivo* modeling of biologically relevant molecular pathways in normal development or physiologic processes, or disease states, guided by human observational data [1]. Some of these mouse models are germline knockouts, with possible adaptive changes that may occur in other associated genes or epigenetic factors, while others conditionally alter gene expression across all or different subsets of vascular or lymphatic endothelial cells. Cre recombinase, or tetracycline-controlled transactivator protein tTA are standard means to achieve conditional gene knockout or knock in, with a goal to better understand the role(s) of specific genes in murine vascular biology with human implications.

Cre recombinase is a 343 amino acid, 38 kDa bacteriophage P1 protein that catalyzes a site specific recombination reaction between two 34-base pair (bp) “locus of crossing over in P1” DNA sequences (called loxP) sites that are absent from the mammalian genome [2–7]. A gene containing two loxP sites flanking essential exons can be deleted in cell types that selectively express Cre recombinase driven by a specific promoter constitutively or inducibly [5–8]. Conditional transgene activation is achieved by placing a flanked loxP (floxed) stop cassette between the promoter and its gene that is removed upon Cre-mediated excision, results in transgene expression [5–7, 9].

Many of these mouse strains alter gene expression in both microvascular and macrovascular endothelial cells in multiple or specific organs, with some also demonstrating altered gene expression in non-endothelial cells such as in hematopoietic stem cells, yolk sac, placenta, osteoblasts, megakaryocytes, hepatocytes, macrophages, lymphocytes, adipocytes, keratinocytes and skeletal muscles, to mention a few [1].

Due to phenotypic and functional differences between microvascular and macrovascular endothelial cells in the same tissues and organs [10–12], developing mouse strains that facilitate selective microvascular endothelial cell gene alternation is important. Such models are relevant to decipher key molecular determinants and signaling pathways in tight junction-forming microvascular endothelial cells that form the blood-brain, blood-nerve, blood-retina and blood-testis barriers. The most commonly used vascular endothelial cell models, based on tyrosine kinase with immunoglobulin-like and EGF-like domains 1 (Tie-1) or TEK receptor tyrosine kinase (Tek, also known Tie-2) alter gene expression in all endothelial cells [1, 13–17]. Data from the mouse sciatic nerve transcriptome demonstrated more diffuse *Tie*- and *Tek* cellular transcript expression in adult (60 day old) mice by single cell whole exome shortgun sequencing (RNA Seq) [18].

The von Willebrand factor (vWF) gene encodes a large glycoprotein transcribed and translated by only microvascular endothelial cells and megakaryocytes that forms multimeric complexes required for platelet adhesion at sites of vascular injury. vWF is stored in Weibel-Palade bodies in endothelial cells and α-granules in megakaryocytes and their derived platelets [19]. vWF is predicted to be involved in chaperone, identical protein and integrin binding activity and implicated in liver and placenta development. A Cre recombinase transgenic mouse model designed to specifically knockout genes in murine brain endothelial cells was developed and published over 10 years ago, driven by a human microvascular endothelial cell-specific promoter (Tg (VWF-Cre)^1304Roho^) [5]; however, these mice are not commercially available or maintained in a known mutant mouse repository for scientific use, questioning strain adequacy or viability. In order to study organ- and time-dependent microvascular endothelial cell signaling pathways in development, post-natal health, aging and disease, we developed a Tamoxifen-inducible vWF Cre recombinase transgenic mouse strain with a structurally stable transgene and highly efficient Cre-mediated recombination, and show restricted microvascular endothelial cell expression in multiple organs, including the brain and sciatic nerves, using validated fluorescent reporter mouse strains.

## Materials and Methods

### Development of Tamoxifen inducible von Willebrand Factor Cre recombinase transgenic mice

The University of Michigan Transgenic Animal Model Core synthesized of a human vWF genomic DNA fragment containing the 734 bp microvascular endothelial cell-specific sequence [20, 21] to drive Cre recombinase fused to a mutant estrogen ligand-binding domain [ERT2] that requires Tamoxifen for activity (CreERT2) followed by a polyadenylation (polyA) signal. The synthesized genomic fragment was subcloned into a CreERT2 plasmid [22] that placed CreERT2 under the control of the synthesized human vWF promoter, and the cloned transgene was verified by DNA sequencing. The cloned transgene was subsequently removed from the plasmid cloning vector and purified for subsequent microinjection. The purified transgene containing the human vWF promoter, inducible Cre (iCre) and polyA signal sequences was microinjected into fertilized mouse eggs obtained by mating mixed strain C57BL/6 X SJL F1 male and female mice, as previously described [23], and transferred to pseudopregnant recipients to produce hemizygous transgenic (designated as vWF-iCre/+) N1 founder mice at the University of Michigan. All hemizygous vWF-iCre/+ mice were transferred to the University of Alabama at Birmingham (UAB) after confirmatory tail snip genotyping.

### Identification of inducible von Willebrand Factor Cre recombinase transgene

All experimental mice were housed in the Research Services Building animal facility, UAB (AAALAC International approved). The studies described were conducted in accordance with all provisions of the Animal Welfare Act, the U.S. Government Principles Regarding the Care and Use of Animals, the Guide (Guide for the Care and Use of Laboratory Animals), National Research Council, Institute of Laboratory Animal Resources, the PHS Policy on Humane Care and Use of Laboratory Animals and the additional local policies of the Center of Comparative Medicine, UAB under approved Institutional Animal Care and Use Committee Protocol # 141009960. Mice were adequately fed with irradiated chow and water provided *ad lib* and housed under micro-isolated, pathogen-free ventilated cages with a 12-hour light-dark cycle compliant with National Institute of Health (NIH) guidelines on animal care. Cages are changed in laminar flow hoods using aseptic micro-isolator techniques, with standard animal husbandry performed by certified veterinary technicians and research staff. All experiments performed were in compliance with the ARRIVE guidelines.

Tail snip genotyping was performed to identify hemizygous mixed strain C57BL/6-SJL vWF- iCre/+ N1 founder mice at 2-3 weeks of age. DNA was extracted using the Viagen Biotech DirectPCR Lysis reagent (catalog # 102-T) followed by DNA purification using isopropanol-ethanol, according to the manufacturer’s instructions. Isolated DNA dissolved in ddH_2_O was quantified and purity verified by absorbance at 260 nm and 260/280 nm absorbance ratios using a Thermo Scientific Nanodrop ND-1000 spectrophotometer with the NanoDrop 1000 3.8.1 software program, and reconstituted at a concentration of 50 ng/µL with sterile ddH_2_O prior to storage at -20°C.

PCR amplification was performed using primers designed to detect both the human vWF promoter and the iCre sequence as follows: vWF-iCre Forward Primer 5’- AATCTTTTCTCCTGCTTTAAAGAAATGTT-3’ and vWF-iCre Reverse Primer 5’- ATCTGTGACAGTTCTCCATCAGGGATCT-3’. Primers were produced by the UAB Heflin Center Genomics Core/ Integrated DNA technologies (www.idtdna.com). PCR amplification was performed by adding 1.0 µL purified tail snip genomic DNA to a mixture of 0.5 µL each of forward and reverse primers from a 10 µM stock concentration with 12.5 µL of 2X Promega GoTaq^®^ Green Master Mix (commercially available premixed ready-to-use solution containing bacterially derived Taq DNA polymerase, dNTPs, MgCl_2_ and reaction buffers at optimal concentrations, catalog # PRM5123) and 10.5 µL of sterile ddH_2_O, with a total of 25 µL PCR reaction mixture. DNA amplification was performed using a Techne TC-412 Thermal cycler programmed as follows: Initiation/Melting 95°C for 5 minutes, 35 cycles for denaturation at 95°C for 30 seconds, annealing at 55°C for 30 seconds and elongation at 72°C for 30 seconds, followed by amplification at 72°C for 5 minutes and termination at 4°C.

PCR products were loaded on a 2% agarose gel in 1X TAE buffer (40 mM **T**ris base, 20 mM **A**cetic acid, and 1 mM **E**thylenediaminetetraacetic acid [EDTA]) containing 0.01% MilliporeSigma™ GelRed™ Nucleic Acid Stain (Fisher Scientific, catalog # SCT123) with electrophoresis performed at 200V for 30 minutes to detect the 604 bp transgene amplicon. GeneRuler 100 bp DNA Ladder (ThermoFisher Scientific, catalog # FERSM0242) was used to confirm band size. Direct or reverse digital images were generated using an AlphaImager HP image documentation system (Cell Biosciences) attached to a Sony ICX267AL 1.39 Megapixel CCD camera and processed using the AlphaView software program. This genotyping protocol was subsequently used to detect the vWF-iCre transgene from the tail snips of all mice when weaned at 3-4 weeks of age.

### Verification of restricted expression using mT/mG mice and determination of Cre recombinase efficiency in the bone marrow

In order to verify that Cre recombinase expression was restricted to vWF-expressing cells (megakaryocytes and microvascular endothelial cells), the 8 week old male N1 founder hemizygous C57BL/6-SJL vWF-iCre/+ mouse was mated with 8 week old female homozygous B6.129(Cg)-Gt(ROSA)26Sor^tm4(ACTB-tdTomato,-EGFP)Luo^/J mice (Jax mice Stock # 007676, kind gift from Dr. Jennifer Pollock, UAB), designated as mT/mG^flox/flox^. This is a mouse strain with a cell membrane-targeted, two-color fluorescent Cre recombinase reporter allele. Prior to Cre recombination with excision of loxP sites, the β-actin promoter driven cell membrane-localized tandem dimer Tomato (mT) red fluorescence protein expression is widely expressed in cells/tissues. The cell membrane-localized green fluorescent protein (mG) replaces mT in Cre-expressing cells/ tissues after recombination and excision [24, 25]. Genotyping was performed to detect the 128 bp mutant and 212 bp wildtype alleles according to the vendor’s protocol (https://www.jax.org/Protocol?stockNumber=007676&protocolID=20368).

Three male and 2 female heterozyous mT/mG (mT/mG^flox/+^) with (vWF-iCre/+) or without (+/+) the hemizygous transgene were injected with 100 mg/kg Tamoxifen (Sigma-Aldrich catalog # T5648) in corn oil (Sigma-Aldrich catalog # C8267) i.p. for 5 consecutive days at 5 weeks of age to evaluate specificity and efficiency of Cre-mediated recombination 3 weeks after injection [26]. Bone marrow was isolated from the femurs and tibia of Tamoxifen-treated mT/mG^flox/+^; vWF- iCre/+ and mT/mG^flox/+^; +/+ mice immediately after euthanasia by cervical disarticulation under deep ketamine/xylazine anesthesia (100 mg/kg/10 mg/kg i.p.) [27]. Four long bones were harvested per mouse and kept on ice prior to bulk processing.

Unfixed bone marrow smears were diffusely spread on Fisherbrand™ Superfrost^TM^ glass slides (Fisher Scientific, catalog # 12-550-14G) without heparin treatment in the dark, carefully stained with 0.45 µM 4′,6-diamidino-2-phenylindole (DAPI) in 1X phosphate buffered saline (PBS) for 5 minutes, washed once with 1X PBS for 5 minutes, and mounted using ProLong^TM^ Gold Antifade aqueous mounting medium (ThermoFisher Scientific, catalog # P36930) with coverslips placed over the entire smear and sealed with nail polish. Slides were immediately visualized using a Nikon ECLIPSE Ci upright epifluorescent microscope attached to a Nikon DS-Qi2 monochromatic camera. Digital photomicrographs generated at 400X magnification to visualize bone marrow cells and detect megakaryocytes throughout each smear using the Nikon NIS Elements AR software program. The efficiency for Cre-mediated recombination was determined in each Tamoxifen-treated mT/mG^flox/+^ mouse as follows: (total number of mG-expressing megakaryocytes divided by total number of mG or mT-expressing megakaryocytes) x 100%, with the mean recombination efficiency calculated.

### Sciatic nerve indirect fluorescent immunohistochemistry reporter protein expression

Sciatic nerves were harvested from both hind limbs prior to removal of femurs and tibia in Tamoxifen-treated mT/mG^flox/+^; vWF-iCre/+ and mT/mG^flox/+^; +/+ mice as described above. Unfixed nerves were immediately embedded in Scigen Tissue Plus^TM^ Optimum Cutting Temperature^®^ (OCT) Compound (Fisher Scientific, catalog # 23-730-571) and stored at -80°C until processed. 10 µm axial cryostat sections were placed on Fisherbrand™ Superfrost™ slides (Fisher Scientific, catalog # 12-550-14G), air dried for 20 minutes at room temperature (RT), fixed with 500 µL of 4% paraformaldehyde in 1X PBS for 30 minutes at RT and washed twice with 1X PBS for 10 minutes. Sections were blocked with 500 µL of 10% normal goat serum (NGS) in 1X PBS, followed by incubation with 1:100 dilution (5 µg/mL) of rabblt anti-red fluorescent protein (RFP) polyclonal IgG antibody (Ab62341) in 2% NGS in 1X PBS at RT for 1 hour. After washing with 2% NGS in 1X PBS for 10 minutes, sections were incubated with 1:500 dilution (4 µg/mL) of goat anti-rabbit IgG (H+L) AlexaFluor^®^594 antibody (Invitrogen, catalog # A-32740) in 2% NGS in 1X PBS for one hour in the dark, with all subsequent processing performed in the dark to prevent photobleaching.

After washing with 2% NGS in 1X PBS for 10 minutes, sections were incubated with 1:500 dilution (10 µg/mL) of rabbit anti-green fluorescent protein (GFP) polyclonal IgG antibody (Ab290) in 2% NGS in 1X PBS for 1 hour, followed by a wash as previously described, then incubated with 1:500 (4 µg/mL) dilution goat anti-rabbit IgG (H+L) AlexaFlour^®^488 antibody (Invitrogen, catalog # A-11008) in 2% NGS in 1X PBS for 1 hour. After washing, sections were stained with 0.45 µM DAPI in 1X PBS for 5 minutes, washed once with 1X PBS for 5 minutes, mounted using ProLong^TM^ Gold Antifade aqueous mounting medium (ThermoFisher Scientific, catalog # P36930) and coverslips placed and sealed with nail polish. Slides were stored at 4°C overnight and visualized with Nikon ECLIPSE Ci upright epifluorescent microscope attached to a Nikon DS-Qi2 monochromatic camera. Digital photomicrographs were generated using the auto-exposure feature (20 to < 1000 ms for the ultraviolet channel and 1s for the red and green channels) to acquire digital images with “look up table” (LUT) adjustments made to reflect observed signal intensities during microscopy in a non-destructive manner during image processing. This takes into account variations in background signal and fluorescent intensities in different specimens. Images were merged using the Nikon NIS Elements AS software program.

### Backcross to SJL using speed congenics

In order to study the role of microvascular endothelial cell-specific gene deletion in murine models of peripheral nerve inflammation (which are relatively mild in C57BL/6 mice), mixed strain hemizygous C57BL/6-SJL vWF-iCre/+ mice were backcrossed to the SJL background by speed congenics, performed at Charles River using their proprietary Marker-Assisted Accelerated Backcrossing (MAX-BAX^®^) protocol. Briefly, a 384 single nucleotide polymorphism (SNP) panel was employed to generate mouse tail snip allelic profiles that were spread across the genome at 7 Mbp intervals for comparison to the wildtype SJL strain reference profile. Each SNP is dimorphic with one allele identified with fluorescent dye 6-Fluorescein amidite (FAM)- labeled complementary oligonucleotide (TaqMan) probe and the other allele detected using fluorescent dye modified *Aequorea victoria* Green Fluorescent Protein (VIC)-labeled probe. Homozygosity at each SNP markers was indicative of genetic alignment with SJL mice. The Percent Match of each mouse with the SJL background was determined by comparing the genotype at each SNP marker, with >99.9% match indicative of complete backcross.

To confirm Tamoxifen-induced specific Cre recombinase expression in microvascular endothelial cells in multiple organs, with higher expression expected relative to megakaryocytes in vWF-iCre/+ SJL mice, 6-9 week old adult male and female mice were mated with age-matched homozygous B6.Cg-*Gt(ROSA)26Sor^tm14(CAG-tdTomato)Hze^*/J (Jax Mice Stock #007914, kind gift from Dr. Karen Gamble, UAB), designated as Ai14^flox/flox^. Prior to Cre-mediated recombination, expression of the CAG promoter driven tandem dimer Tomato (tdTomato [tdT]) red fluorescent protein is prevented by a loxP-flanked stop cassette. Following Cre recombinase excision, transcription with subsequent translation of robust red fluorescent protein expression occurs in Cre-expressing cells/ tissues [28]. Genotyping was performed to detect the 196 bp mutant and 297 bp. wildtype alleles according to the vendor’s protocol (https://www.jax.org/Protocol?stockNumber=007914&protocolID=29436). Heterozygous mixed strain C57BL/6-SJL Ai14^flox/+^; vWF-iCre/+ F1 mice were intercrossed with homozygous C57BL/6 Ai14^flox/flox^ mice to generate mixed strain C57BL/6-SJL Ai14^flox/flox^; vWF-iCre/+ F2 mice for subsequent reporter expression evaluation.

Cohorts of 5-7 week old mixed strain C57BL/6-SJL Ai14^flox/flox^; vWF-iCre/+ F2 mice were injected with 100 mg/kg Tamoxifen (Sigma-Aldrich catalog # T5648) in corn oil (Sigma-Aldrich catalog # C8267) i.p. for 5 consecutive days to evaluate specificity of Tamoxifen-induced Cre recombination, with mixed strain C57BL/6-SJL Ai14^flox/flox^; vWF-iCre/+ injected with equivalent volumes of corn oil serving as controls for Tamoxifen-independent recombination (which has been described in Cre expressing Ai14^flox/flox^ mice)[6, 7]. Mixed strain C57BL/6-SJL Ai14^flox/flox^; +/+ were also injected with 100 mg/kg Tamoxifen (Sigma-Aldrich catalog # T5648) in corn oil (Sigma-Aldrich catalog # C8267) for 5 consecutive days or corn oil alone to serve as controls for non-specific or cell-independent red fluorescent protein expression. 24-25 days after injection, mice underwent whole body perfusion fixation via intracardiac infusion of 4% paraformaldehyde in 1X PBS at 37°C under deep ketamine/ xylazine (100 mg/kg/ 10 mg/kg) anesthesia after a sterile 4°C 1X PBS infusion flush, as published [29, 30] (courtesy of Dr. Scott Wilson, UAB).

The femur and tibial bones on both sides were harvested and processed for bone marrow smear analyses in the dark on the day of tissue harvest, as previously described. Both sciatic nerves, brain, spleen, kidneys and gastrocnemius muscles were harvested from each mouse, additionally fixed for 24 hours by immersion in 4% paraformaldehyde in 1X PBS at 4°C, washed three times in 1X PBS for 10 minutes and cryopreserved for 12-15 hours in 30% sucrose weight/volume in 1X PBS, washed as described above, and embedded in OCT Compound with storage at -80°C until further processing.

### Direct fluorescent immunohistochemistry to evaluate selective microvascular endothelial cell Cre expression in reporter mice

Paraformaldehyde-fixed, sucrose cryoprotected frozen sciatic nerves, brain, spleen, kidneys and gastrocnemius muscles were evaluated to determine specific Tamoxifen-inducible Cre recombinase expression in microvascular endothelial cells in the previously described cohort of mixed strain C57BL/6-SJL Ai14^flox/flox^; vWF-iCre/+ F2 mice and littermate controls injected with corn oil alone or Tamoxifen-injected Ai14^flox/flox^; +/+ F2 mice.

10 µm axial or longitudinal cryostat sections were placed on Fisherbrand™ ColorFrost™ Plus slides (Fisher Scientific, catalog # 12-550-18) and air dried for 12-15 hours overnight in the dark at RT. Subsequent processing was performed in the dark to prevent endogenous tdT red fluorescence photobleaching. Sections were incubated with 500-1000 µL of 1:200 dilution (5 µg/mL) of *Lycoperiscon esculentum* agglutinin lectin-DyLight^®^ 488 (fluorochrome-conjugated Tomato lectin [TL], which is the most sensitive marker for mouse vascular endothelial cells, binding to N-acetylglucosamine residues [31]) in 1X PBS at RT for 1 hour. Following 1-2 washes with 1X PBS for 10 minutes each, autofluorescence quenching was performed using Vector^®^ TrueVIEW^®^ Autofluorescence Quenching Kit (Vector Laboratories, catalog # SP-8400) at RT, according to the vendor’s protocol.

Following this process, sections were washed with 1X PBS 1-2 times for 10 minutes each, and treated with 500-1000 µL of 0.45 µM DAPI in 1X PBS for 5 minutes at RT. Following washes as described above, sections were mounted with VECTASHIELD Vibrance^®^ Antifade Medium (provided with autofluorescence quenching kit) with coverslips placed over slides and sealed with nail polish. Slides were stored in the dark at 4°C for no longer than 48 hours, and viewed using a Nikon ECLIPSE Ci upright epifluorescent microscope attached to a Nikon DS-Qi2 monochromatic camera. Digital photomicrographs were generated and merged using the Nikon NIS Elements AR software program as previously described. To further verify reporter expression on microvascular endothelial cells and support the utility of TL as a sensitive mouse vascular endothelial cell marker, CD31 indirect fluorescent immunohistochemistry (clone 390, ThermoFisher Scientific catalog # 14-0311-82) was performed on sciatic nerve axial sections from the same Tamoxifen-injected Ai14^flox/flox^; vWF-iCre/+ SJL F2 mice, as previously published [32].

To determine whether the Tamoxifen-inducible Cre recombinase is expressed by lymphatic endothelial cells, longitudinal sections of spleen and kidney sections from the previously described cohort of Tamoxifen-injected mixed strain C57BL/6-SJL Ai14^flox/flox^; vWF-iCre/+ F2 mice were sectioned, mounted, air-dried in the dark and blocked with 10% NGS in 1X PBS as described above. Sections were incubated with 500-1000 µL of 1:200 dilution (2.5 µg/mL) of hamster anti-mouse Podoplanin IgG antibody (clone eBio8.1.1 (8.1.1); ThermoFisher Scientific catalog # 14-5381-82) diluted in 2% NGS in 1X PBS and incubated overnight at 4°C in the dark. Following washes with 2% NGS in 1X PBS, sections were treated with 500-1000 µL of 1:500 dilution (4 µg/mL) of goat anti-hamster IgG (H+L) AlexaFluor^®^ 488 antibody (ThermoFisher Scientific catalog # A78963) diluted in 2% NGS in 1X PBS and incubated for 1 hour at RT in the dark. Sections were washed, stained with 0.45 µM DAPI in 1X PBS, mounted with Vectashield and viewed, imaged and processed using Nikon ECLIPSE Ci upright epifluorescent microscope, DS-Qi2 monochromatic camera and NIS Elements AS software program, as described above.

Paraformaldehyde-fixed bone marrow smears were generated on the day of whole animal fixation to verify retained megakaryocyte Cre expression in mixed strain C57BL/6-SJL Ai14^flox/flox^; vWF-iCre/+ F2 mice compared to littermate controls injected with corn oil alone or Ai14^flox/flox^; +/+ F2 mice in the dark , as previously described. Splenic megakaryocytes were identified in paraformaldehyde-fixed, sucrose cryoprotected cryostat sections primarily processed to detect Cre expression in microvascular endothelial cells, as described above, with comparative levels of expression qualitatively determined based on red fluorescence mean fluorescent intensity. Slides were viewed at 400X magnification using a Nikon ECLIPSE Ci upright epifluorescent microscope attached to a Nikon DS-Qi2 monochromatic camera, with digital photomicrographs generated and merged as previously described.

### Transgene copy number and chromosomal insertion analysis

The vWF-iCre transgene copy analysis was performed in hemizygous vWF-iCre/+ SJL mice (formally designated as SJL.Cg-Tg(VWF-Cre/ERT2)C1014/UbeeMmmh; Research Resource Identifier [RRID]:MMRRC_071293-MU) by the University of Missouri MMRRC using their validated proprietary protocols adapted from a prior publication [33]. Transgene integrity and chromosomal integration site analyses were performed by Cergentis B.V. (Utrecht, Netherlands) on viable cryopreserved splenocytes from a male adult vWF-iCre/+ SJL mouse using their recommended splenocyte isolation and proprietary Target Locus Amplification (TLA) analysis protocols [34].

## Results

Pictorial representation of the vWF-iCre DNA construct and representative mouse tail snip DNA genotyping identifying the 604 bp transgene is shown in **Figure 1**. The complete vWF-iCre transgene sequence containing the human vWF promoter with the microvascular endothelial cell-selective sequence (position -487 to +246 of the human vWF gene) [20, 21] located at transgene position 236 to 969 is shown in **Supplementary File 1**. Following microinjection into fertilized mouse eggs obtained by mating mixed strain C57BL/6 X SJL F1 male and female mice, and implantation into pseudopregnant females, 143 pups (77 males and 66 females) were born. Tail snip genotyping identified 7 male and 6 female mixed strain C57BL/6-SJL hemizygous vWF-iCre/+ founder mice. 1 male founder mouse (with highest SJL background) was mated with two mT/mG^flox/flox^ female mice to generate mT/mG^flox/+^; vWF-iCre/+ mice to evaluate Tamoxifen-dependent Cre recombination specificity and efficiency. Representative tail snip genotyping data are shown in **Supplementary Figure 1A**.

**Figure 1.**
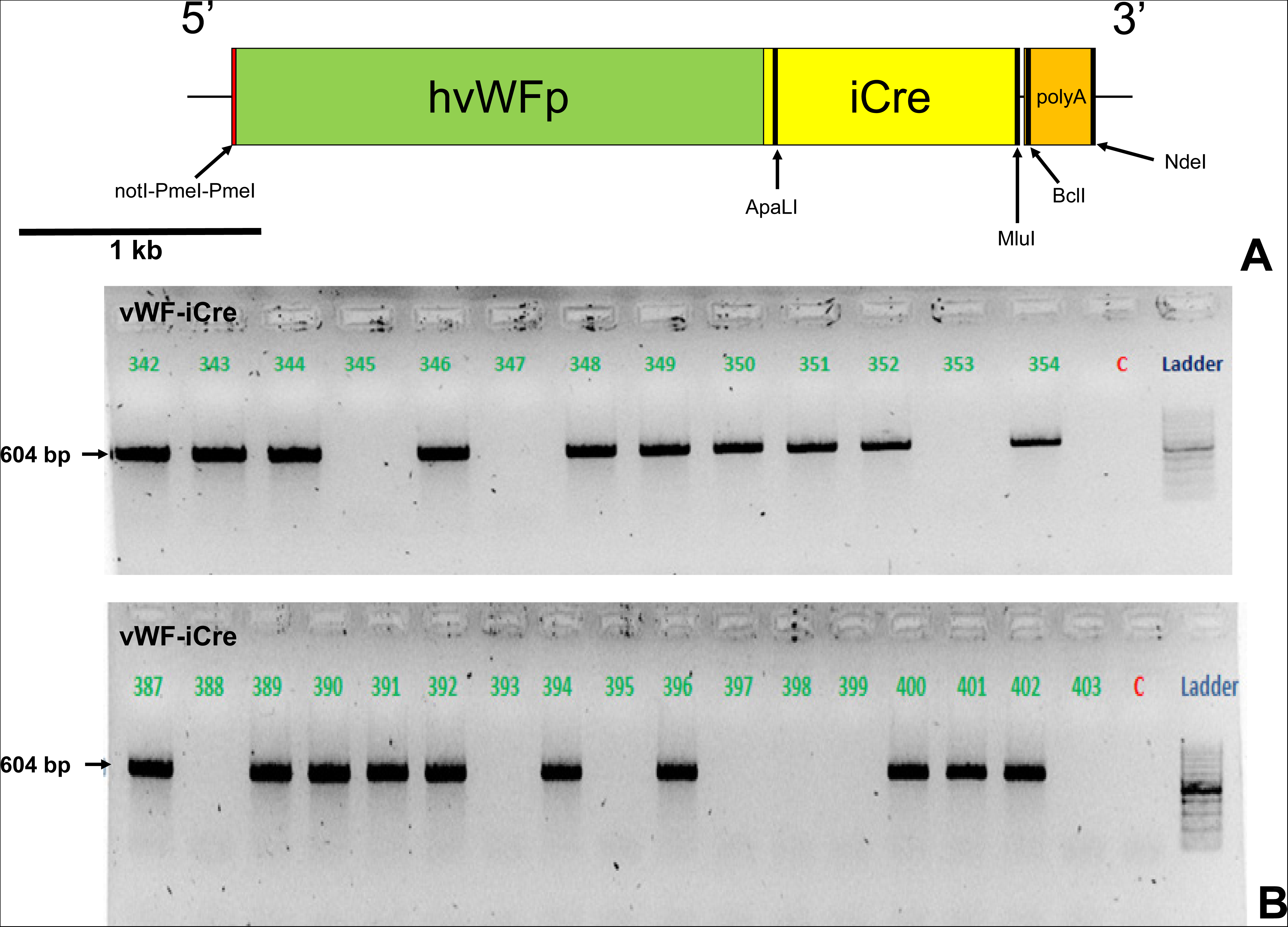
vWF-iCre DNA construct and representative genotyping. Pictorial representation of the vWF-iCre transgene is shown, consisting of the microvascular endothelial cell-selective human von Willebrand factor promoter (hvWFp), Cre recombinase fused to a mutant estrogen ligand-binding domain, CreERT2 (iCre), polyadenylation (polyA) and restriction endonuclease (notI, PmeI, ApaLI, MluI, BclI and NdeI) sequences (**A**). Reverse digital photomicrographs of 2% agarose electrophoresis gels in 1X TAE buffer from a cohort of mice show representative tail snip genotyping results, with the vWF-iCre transgene identified at 604 bp (**B**). C represents lanes with absent DNA (ddH_2_O) template controls.

Unfixed bone marrow smears from mT/mG^flox/+^; vWF-iCre/+ F1 mice demonstrated megakaryocyte-specific mT to mG expression following Tamoxifen-induced recombination (**Figure 2A-L**) that did not occur in mT/mG^flox/+^; +/+ (**Figure 2M-P**). These data show that Tamoxifen-induced bone marrow megakaryocyte specific Cre expression occurs mT/mG^flox/+^; vWF-iCre/+ mice, supportive of transgene specificity. Mean Cre recombination efficiency was 91.2% (range 75.9-100.0, N=5). Sciatic nerve axial sections from mT/mG^flox/+^; vWF-iCre/+ F1 mice showed endoneurial microvessel-like structure mT to mG expression following Tamoxifen-induced recombination (**Figure 3A-D**) that did not occur in control mT/mG^flox/+^; +/+ mice (**Figure 3E-H**), further supported by higher magnification evaluation of these microvessel-like structures (**Figure 3I-X)**. These data support the notion that the vWF-iCre transgene is specifically expressed in bone marrow megakaryocytes and structures consistent with sciatic nerve endoneurial microvessels, with high efficiency *in vivo* Tamoxifen-driven recombination.

**Figure 2.**
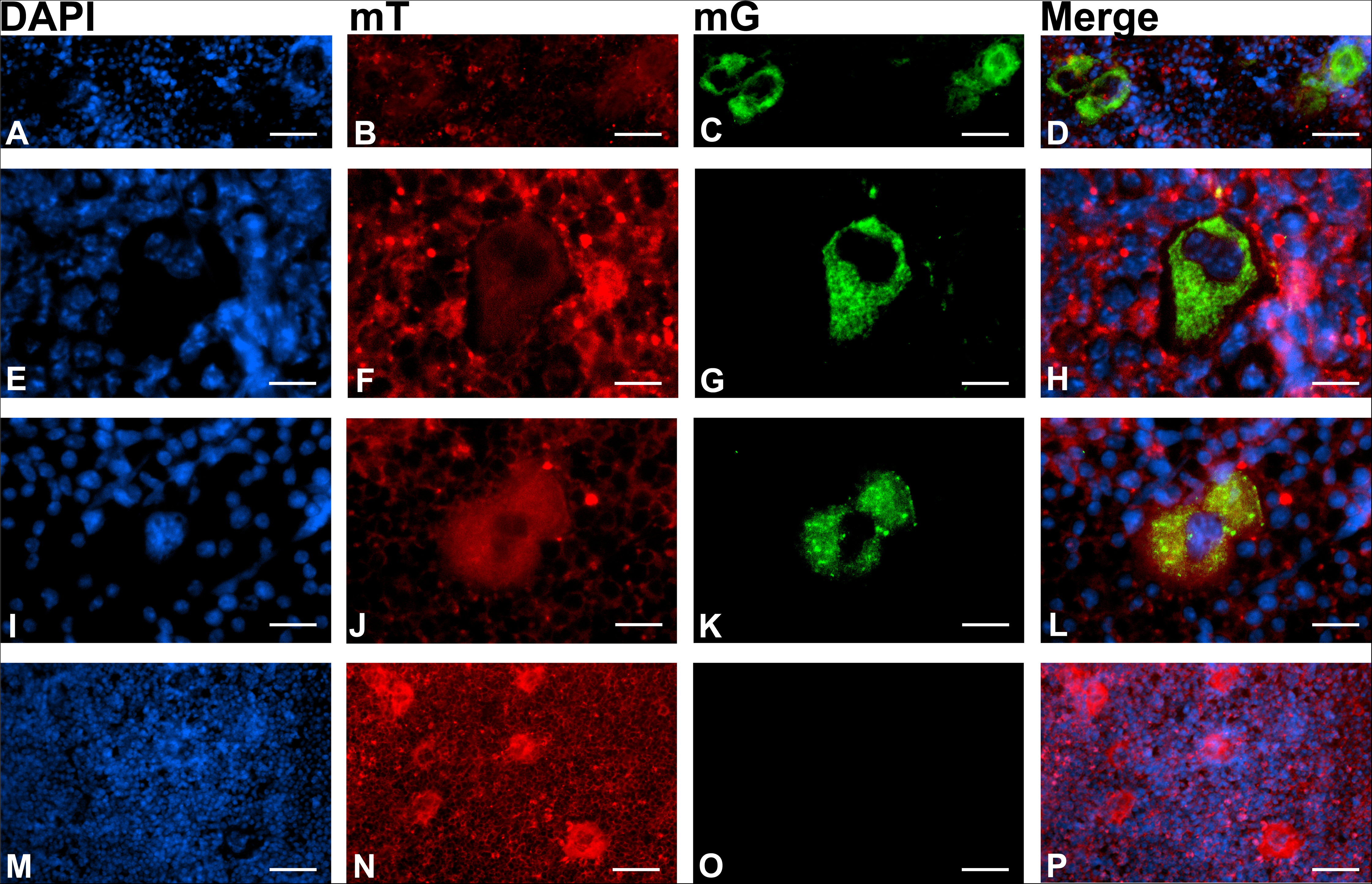
Tamoxifen-induced Cre recombinase expression in bone marrow megakaryocytes of mT/mG^flox/+^; vWF-iCre/+ C57BL/6-SJL mice. Representative digital photomicrographs of unfixed bone marrow smears from mT/mG^flox/+^; vWF-iCre/+ F1 mice show megakaryocyte-specific mT to mG expression following Tamoxifen-induced recombination (**A-L**) that does not occur in control mice that lack the vWF-iCre transgene (**M-P**). Scale bars: A-D and M-P = 100 µm, E-L = 50 µm.

**Figure 3.**
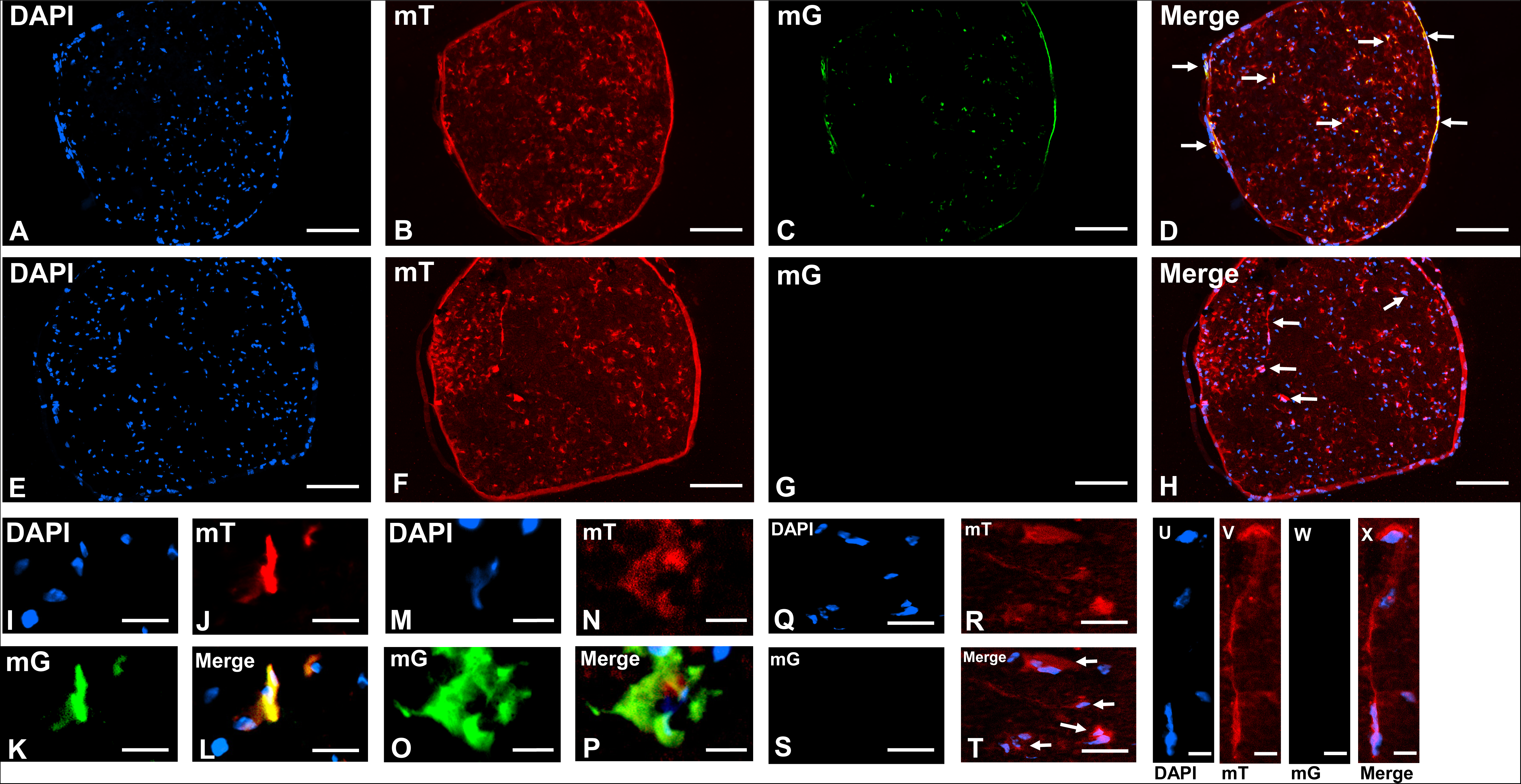
Tamoxifen-induced Cre recombinase expression in sciatic nerve microvessels in mT/mG^flox/+^; vWF-iCre/+ C57BL/6-SJL mice. Representative digital photomicrographs of paraformaldehyde-fixed cryostat axial sciatic nerve sections from mT/mG^flox/+^; vWF-iCre/+ F1 mice show microvessel-like structure mT to mG expression following Tamoxifen-induced recombination, as detected by anti-RFP and anti-GFP antibodies (**A-D**, white arrows) that does not occur in control mice without the vWF-iCre transgene (**E-H**, white arrows). Higher magnification cropped digital photomicrographs of sciatic nerve endoneurial microvessel-like structures demonstrating mT to mG expression to support tissue-specific inducible Cre recombinase expression are shown in **I-P**, with vWF-iCre transgene absent littermate control endoneurial microvessel-like structures without mT to mG expression shown in **Q-X** (white arrows). Scale bars: A-H = 100 µm, I-L and Q-T = 20 µm, M-P and U-X = 5 µm.

The male mixed strain C57BL/6-SJL vWF-iCre/+ founder mouse was mated with wildtype female SJL mice, and offspring genotyped to identify vWF-iCre/+ mice. The Percentage Match, indicative of alignment with wildtype SJL mice, was determined by tail snip genotyping and progeny with highest SJL background were backcrossed with wildtype SJL mice. Congenic SJL vWF-iCre/+ mice were obtained at N6, with identification of a donor genetic sequence on chromosome 1. Subsequent analyses determined that a single copy of the vWF-iCre transgene without structural variants stably integrated at chromosome 1:132,314,865-132,905,344 with a sequence mutation A>T at position 3180 in the iCre region. This was associated with a 125 kb deletion in chromosome 1 involving exons 1-11 of contactin 2 (*Cntn2*) and exons 3-32 of neurofascin (*Nfasc*), as shown in **Figure 4**. Following intercrossing wildtype and vWF-iCre/+ SJL mice for >10 generations, Ai14^flox/flox^ mice were mated with vWF-iCre/+ SJL mice to generate Ai14^flox/flox^; vWF-iCre/+ mice to directly evaluate specific Tamoxifen-induced microvascular endothelial cell Cre expression and evaluate for Tamoxifen-independent and non-specific Cre expression in multiple organs. Representative DNA tail snip genotyping is shown in **Supplementary Figure 1B.**

**Figure 4.**
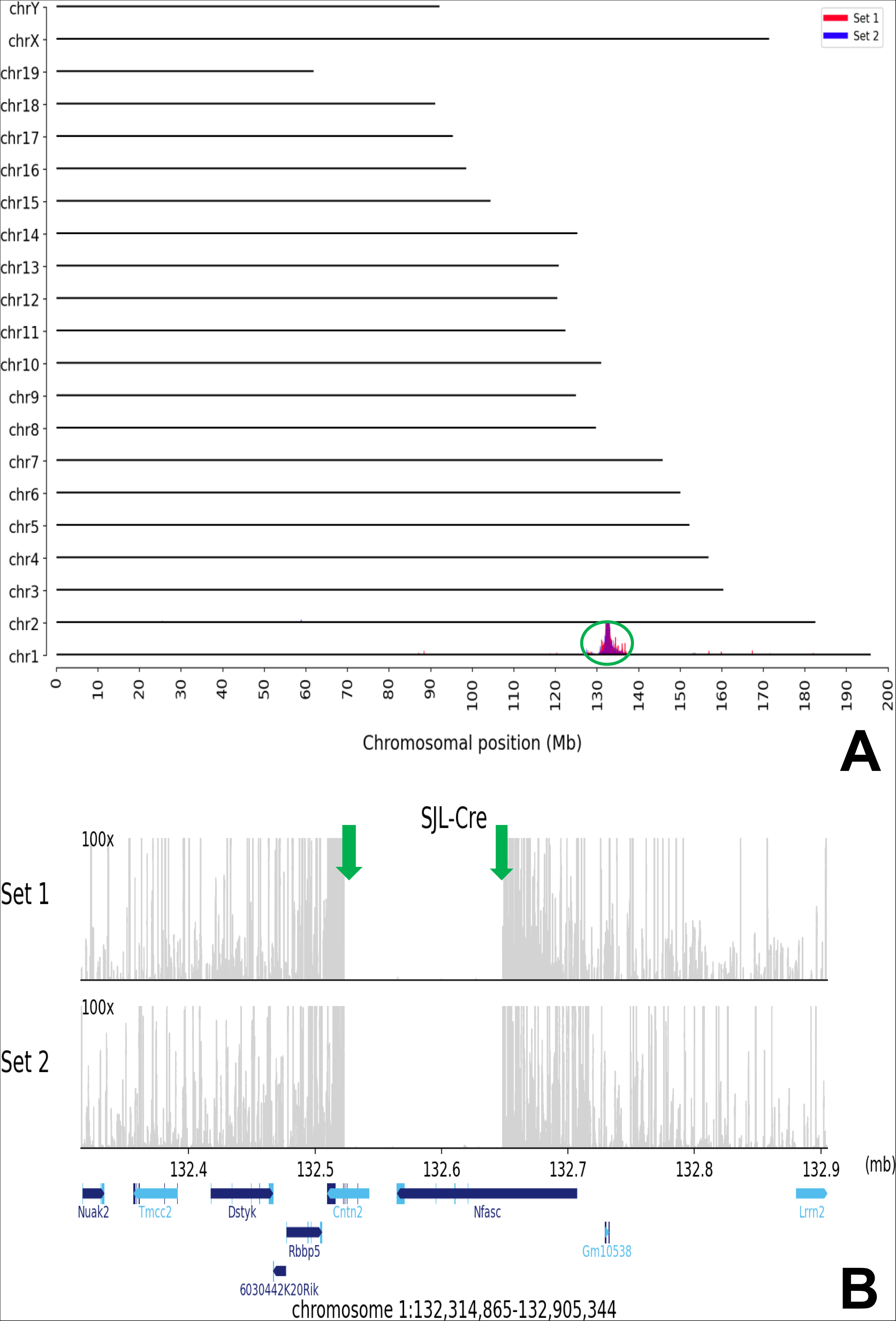
vWF-iCre transgene integration site. TLA was performed using two specific independent primer sets binding to the transgene iCre and bgh-polyA sequences, with next generation sequencing reads aligned to the vector sequence and mouse mm10 reference genome sequence. Whole genome plot using primer set 1 (red) and primer set 2 (blue) demonstrates the transgene integration site on chromosome 1 (**A**). The y-axis represents chromosomes and the x-axis represents chromosomal location. The TLA sequence coverage (grey lines) across the vector integration site (chr1:132,314,865-132,905,344) using both primer sets is shown in **B**, with the green arrows indicative of the location of breakpoint sequence. The 125 kb genomic sequence between the breakpoints has been deleted, as well as vWF-iCre transgene nucleotides at position 1-9 and 3540-3543 that encoded notI, PmeI and NdeI endonucleases. Y-axes are limited to 100x and with light blue and dark blue lines with inscriptions below the x-axis indicating known encoded genes.

Microvascular endothelial cell specific Cre expression was observed in the sciatic nerve endoneurial microvascular endothelial cells of Ai14^flox/flox^; vWF-iCre/+ SJL F2 mice following Tamoxifen-induced recombination based on tdT expression co-localizing with Tomato Lectin (TL) green fluorescence, as shown in **Figure 5A-D**. This is in contrast to absent tdT expression in the sciatic nerves of Ai14^flox/flox^; +/+ SJL F2 control mice (**Figure 5E-H**). Sciatic nerve endoneurial microvessels clearly demonstrated tdT expression in Ai14^flox/flox^; vWF-iCre/+ SJL F2 mice to further support specific Tamoxifen-inducible Cre recombinase expression, as shown in **Figure 5I-X**. Comparative analyses of TL with CD31 immunostaining showed that tdT co- localization was similar on sciatic nerve CD31+ endoneurial microvascular endothelial cells (**Supplementary Figure 2**), supporting the utility of TL to rapidly identify vascular endothelium in reporter mice following *in situ* perfusion fixation [31]. Parenchymal brain microvessels from Tamoxifen-injected Ai14^flox/flox^; vWF-iCre/+ SJL F2 mice also showed tdT co-localizing with TL green fluorescence, consistent with microvascular endothelial cell-specific Cre recombinase expression (**Figure 6A-D**). This is in contrast to absent tdT in the brains of Ai14^flox/flox^; +/+ SJL F2 control mice, as shown in **Figure 6E-H**. Brain microvessels at high magnification further demonstrated tdT expression in Ai14^flox/flox^; vWF-iCre/+ SJL F2 mice compared to absent expression in Ai14^flox/flox^; +/+ SJL F2 control mice, consistent with Tamoxifen-inducible microvascular endothelial cell-specific Cre recombinase expression (**Figure 6I-T**).

**Figure 5.**
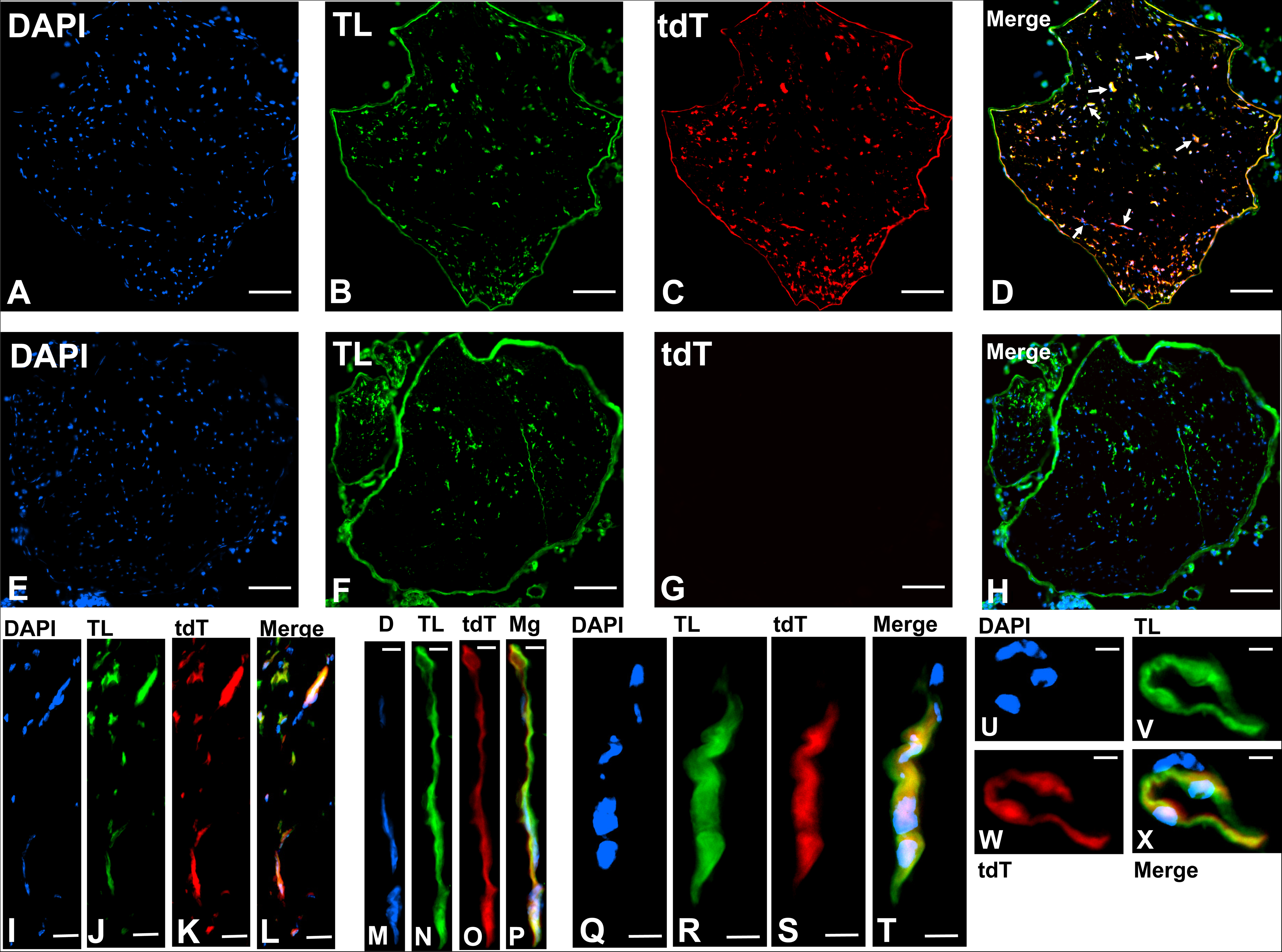
Tamoxifen-induced Cre recombinase expression in sciatic nerve microvascular endothelial cells of Ai14^flox/flox^; vWF-iCre/+ SJL F2 mice. Representative digital photomicrographs of paraformaldehyde-fixed, sucrose cryopreserved cryostat axial sciatic nerve section from an Ai14^flox/flox^; vWF-iCre/+ SJL F2 mouse shows tdT expression co-localizing with Tomato Lectin (TL) green fluorescence, consistent with microvascular endothelial cell-specific Cre recombinase expression following Tamoxifen-induced recombination (**A-D**, white arrows). Cre expression is absent from the sciatic nerve in a control mouse that lacks the vWF-iCre transgene (**E-H**). Higher magnification cropped digital photomicrographs of sciatic nerve endoneurial endothelial cells demonstrating tdT expression to support Tamoxifen-inducible Cre-mediated recombination are shown in **I-X**. Scale bars: A-H = 100 µm, I-K = 50 µm, M-P = 5 µm, Q-T = 20 µm, U-X = 10 µm.

**Figure 6.**
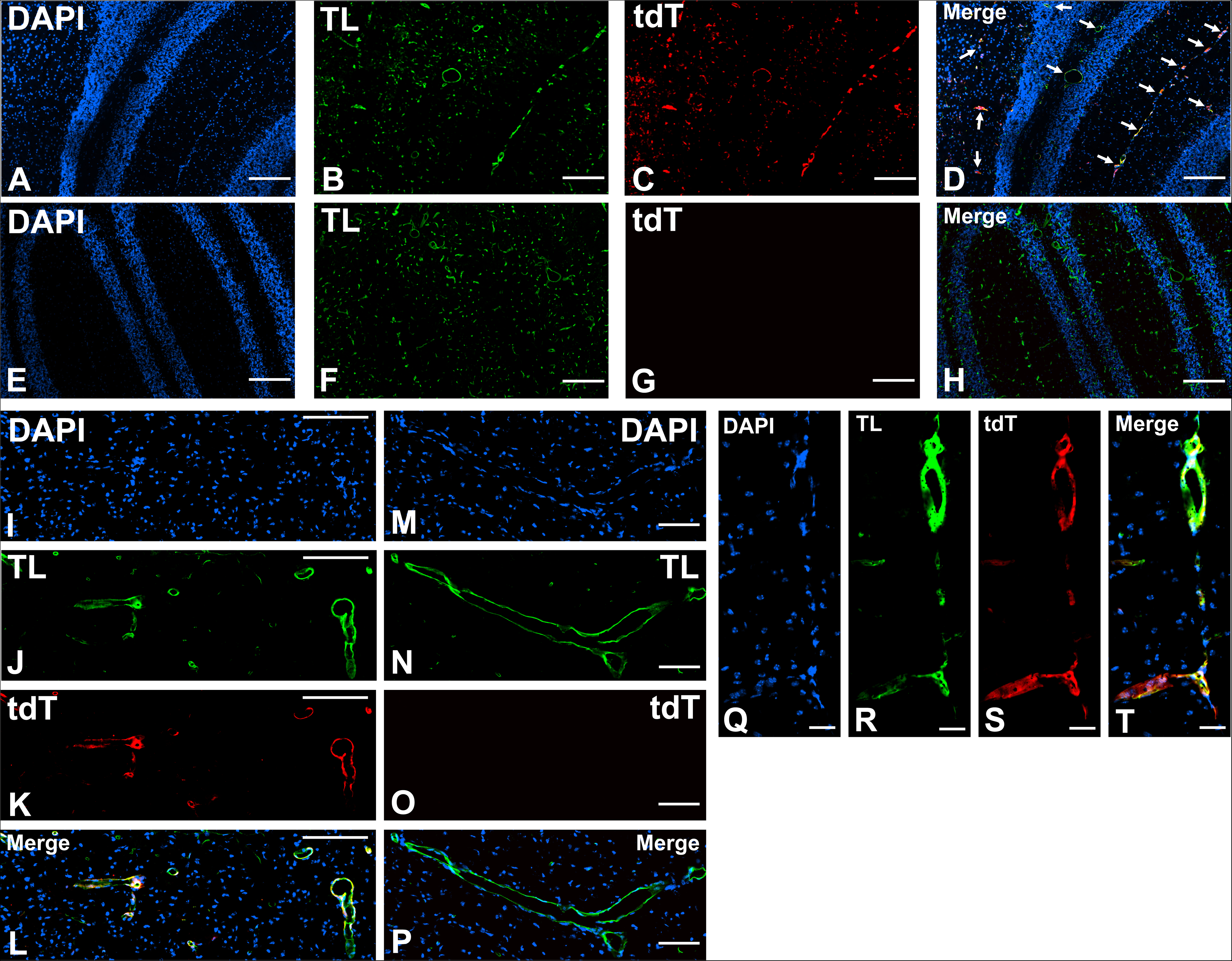
Tamoxifen-induced Cre recombinase expression in brain microvascular endothelial cells of Ai14^flox/flox^; vWF-iCre/+ SJL F2 mice. Representative digital photomicrographs of paraformaldehyde-fixed, sucrose cryopreserved cryostat longitudinal brain section from an Ai14^flox/flox^; vWF-iCre/+ SJL F2 mouse shows tdT expression co-localizing with TL green fluorescence, consistent with microvascular endothelial cell-specific Cre recombinase expression following Tamoxifen-induced recombination (**A-D**, white arrows). Cre expression is absent from the brain of a control mouse without the vWF-iCre transgene (**E-H**). Higher magnification cropped digital photomicrographs of brain microvessels with tdT red fluorescence to support Tamoxifen-inducible Cre recombinase expression are shown in **I-L** compared to absent tdT microvascular endothelial cell expression in the brain of a mouse without the vWF- iCre transgene, shown in **M-P**. Higher magnification brain microvessels with endothelial cell-specific tdT expression following Tamoxifen-induced recombination in an Ai14^flox/flox^; vWF-iCre/+ F2 mouse are also shown (**Q-T**). Scale bars: A-H = 200 µm, I-L = 50 µm, M-T = 25 µm.

Longitudinal sections from Tamoxifen-injected Ai14^flox/flox^; vWF-iCre/+ SJL F2 mouse spleens showed microvascular endothelial cell tdT co-localization, with absent tdT expression in Ai14^flox/flox^; +/+ SJL F2 control mice, in support of microvascular endothelial cell-specific Cre recombinase expression following Tamoxifen-induced recombination (**Figure 7A-I**). TL also stains the structural reticular meshwork of the spleen, as previously described [31]. Low tdT expression was observed in splenic megakaryocytes of Ai14^flox/flox^; vWF-iCre/+ SJL F2 mice (**Figure 7J-K**) compared to the much higher expression in microvascular endothelial cells seen with lower magnification images, consistent with Tamoxifen-induced Cre recombinase expression driven by a microvascular endothelial-cell selective vWF promoter.

**Figure 7.**
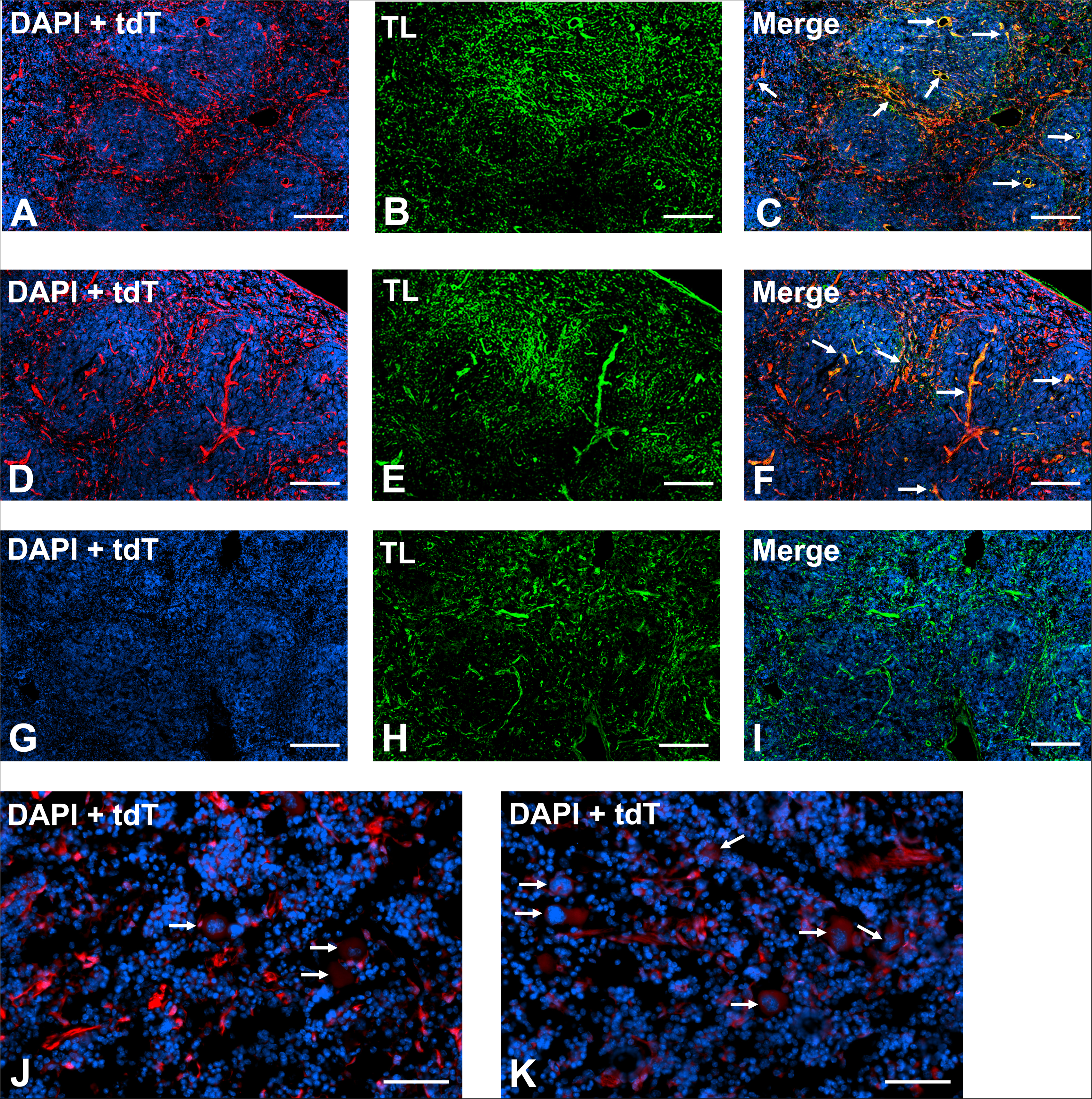
Tamoxifen-induced Cre recombinase expression in spleen microvascular endothelial cells and splenic megakaryocytes of Ai14^flox/flox^; vWF-iCre/+ SJL F2 mice. Representative digital photomicrographs of paraformaldehyde-fixed, sucrose cryopreserved cryostat longitudinal spleen sections from an Ai14^flox/flox^; vWF-iCre/+ SJL F2 mouse shows tdT expression co- localizing with TL green fluorescence, consistent with microvascular endothelial cell-specific Cre recombinase expression following Tamoxifen-induced recombination (**A-F**, white arrows). TL also stains the spleen structural reticular meshwork (background). Cre expression was absent from the spleen of a control mouse lacking the vWF-iCre transgene (**G-I**). Representative digital photomicrographs of paraformaldehyde-fixed, sucrose cryopreserved cryostat longitudinal spleen sections from Ai14^flox/flox^; vWF-iCre/+ SJL F2 mice shows low tdT expression in splenic megakaryocytes (**J-K**, white arrows) compared to high expression in microvascular endothelial cells seen with lower magnification images, consistent with Tamoxifen-induced Cre recombinase expression driven by a selective microvascular endothelial cell vWF promoter. Scale bars: A-C and G-K = 200 µm, D-F = 100 µm.

Longitudinal kidney sections from Ai14^flox/flox^; vWF-iCre/+ SJL F2 mice also showed tdT co- localizing with TL green fluorescence in the cortex following Tamoxifen-induced recombination that was absent in Ai14^flox/flox^; +/+ SJL F2 control mice, consistent with microvascular endothelial cell-specific Cre recombinase expression (**Figure 8A-H**). Medullary microvessels demonstrating tdT expression following Tamoxifen-induced recombination were also observed in Ai14^flox/flox^; vWF-iCre/+ SJL F2 mice (**Figure 8I-L**, white arrows). Ai14^flox/flox^; vWF-iCre/+ SJL F2 mice further demonstrated microvascular endothelial cell tdT expression in glomerular capillaries and post-capillary venules that were absent from peritubular capillaries, as shown in **Figure 8M-P**. This microvascular endothelial cell Tamoxifen-inducible Cre recombinase expression pattern is consistent with known heterogeneous murine renal microvascular endothelial cell vWf expression *in situ* [35]. Microvascular endothelial cell tdT expression was also observed in the gastrocnemius muscles of Ai14^flox/flox^; vWF-iCre/+ SJL F2 mice following Tamoxifen-induced recombination (**Figure 9A-D**). An unexpected myofiber sarcoplasmic tdT mosaic expression was also observed. However, tdT expression was absent from Ai14^flox/flox^; +/+ SJL F2 control mice (**Figure 9E-H**). No murine myogenic basic Helix-Loop-Helix (bHLH) transcription factor (*Myod1*, *Myf5*, *Myf6* and *Myog*) functional sequences [36] were identified in the vWF-iCre transgene sequence (DNA generated from amino acid sequences using Sequence Manipulation Suite Reverse Translate [https://www.bioinformatics.org/sms2/rev_trans.html]). Endomysial microvessels demonstrated endothelial cell tdT expression to further support microvascular endothelial cell-specific Tamoxifen-inducible Cre recombinase expression, as shown in **Figure 9I-L**.

**Figure 8.**
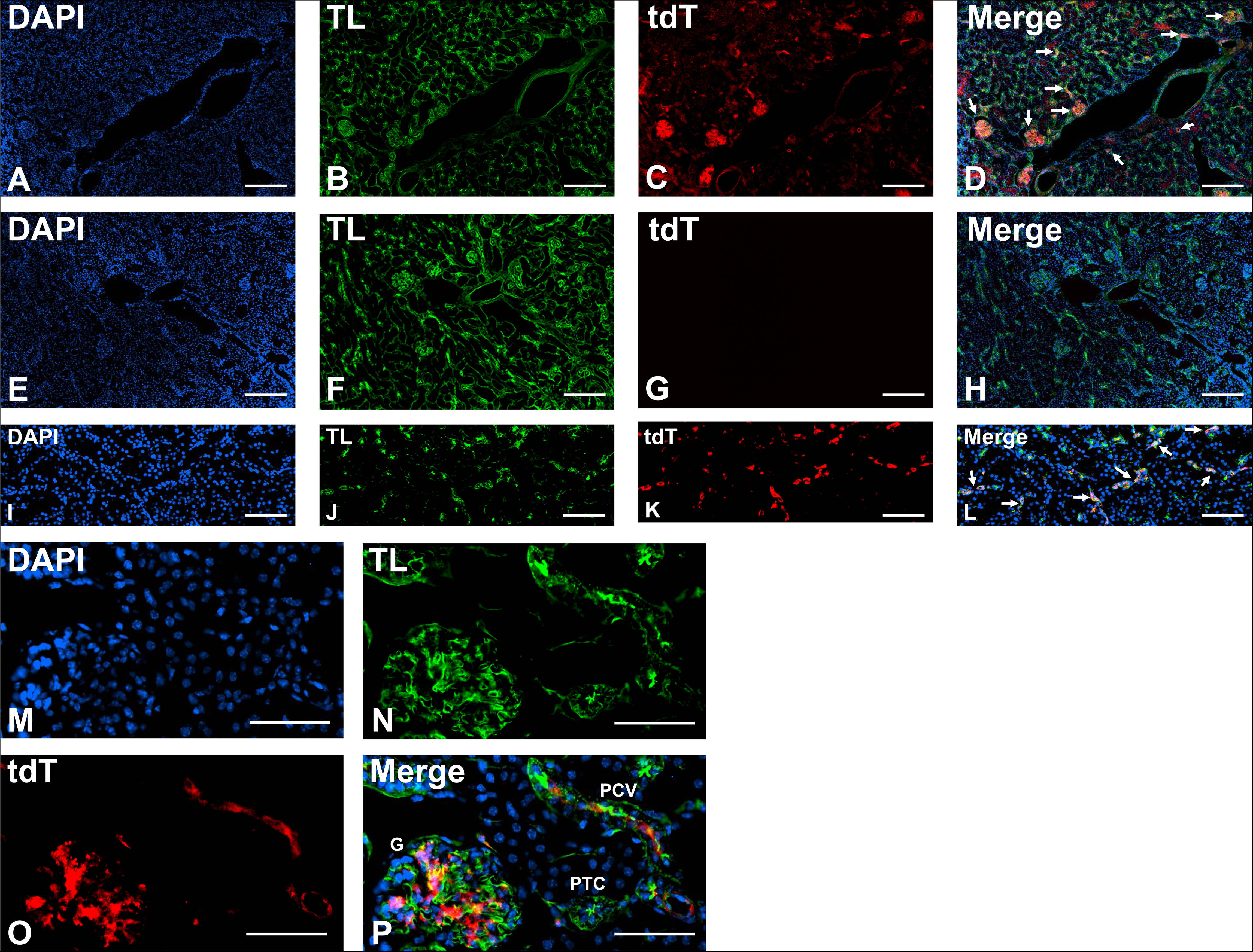
Tamoxifen-induced Cre recombinase expression in kidney microvascular endothelial cells of Ai14^flox/flox^; vWF-iCre/+ SJL F2 mice. Representative digital photomicrographs of paraformaldehyde-fixed, sucrose cryopreserved cryostat longitudinal kidney sections from an Ai14^flox/flox^; vWF-iCre/+ SJL F2 mouse shows tdT expression co-localizing with TL green fluorescence, consistent with microvascular endothelial cell-specific Cre recombinase expression following Tamoxifen-induced recombination in the cortex (**A-D**, white arrows). Cre expression was absent from the cortical microvessels of a control mouse without the vWF-iCre transgene (**E-H**). Medullary microvessels demonstrating tdT expression following Tamoxifen- induced recombination in an Ai14^flox/flox^; vWF-iCre/+ SJL F2 mouse are shown (**I-L**, white arrows). Higher magnification cropped digital photomicrographs of the kidney cortex shows microvascular endothelial cell tdT expression in capillaries within a glomerulus (G) and post- capillary venule (PCV) that is absent from a peritubular capillary (PTC) in an Ai14^flox/flox^; vWF- iCre/+ SJL F2 mouse, supporting selective kidney microvascular endothelial cell Tamoxifen- inducible Cre recombinase expression (**M-P**). Scale bars: A-H = 200 µm, I-L = 100 µm, M-P = 50 µm.

**Figure 9.**
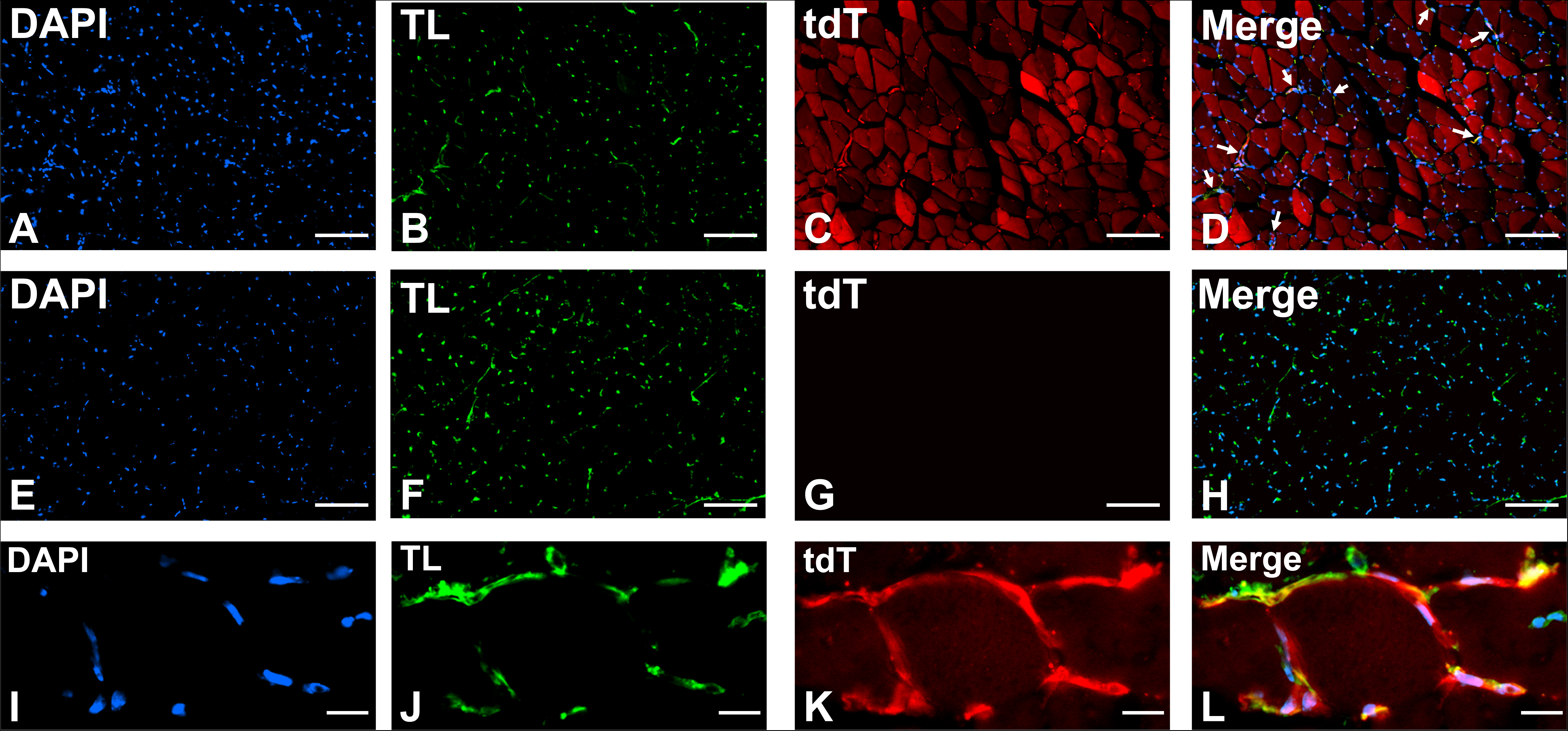
Tamoxifen-induced Cre recombinase expression in gastrocnemius muscle microvascular endothelial cells of Ai14^flox/flox^; vWF-iCre/+ SJL mice. Representative digital photomicrographs of paraformaldehyde-fixed, sucrose cryopreserved cryostat axial muscle sections from an Ai14^flox/flox^; vWF-iCre/+ SJL F2 mouse show tdT expression co-localizing with TL green fluorescence, consistent with microvascular endothelial cell-specific Cre recombinase expression following Tamoxifen-induced recombination (**A-D**, white arrows). A mosaic pattern of tdT expression is unexpectedly observed within muscle fascicles. Cre expression was absent from the gastrocnemius muscles in a control mouse lacking the vWF-iCre transgene (**E-H**). Higher magnification cropped digital photomicrographs of endomysial microvascular endothelial cells demonstrating tdT expression to support Tamoxifen-inducible Cre recombinase expression are shown in **I-L**. Scale bars: A-H = 100 µm, I-L = 10 µm.

Axial sections from Tamoxifen-injected Ai14^flox/flox^; vWF-iCre/+ SJL F2 mouse sciatic nerves showed epineurial arteriole and venule endothelial cell tdT localization (white arrows) in addition to endoneurial microvessel endothelial cell expression (asterisk), as shown in **Figure 10A-D**. The epineurial blood vessels had diameters less than 50 µm. No large arteries or veins were present in the retained epineurium or preserved from the vasa nervonum. Tdt expression was absent from large splenic arteries (**Figure 10E-H**) and veins (**Figure 10I-L**) in Tamoxifen-injected Ai14^flox/flox^; vWF-iCre/+ SJL F2 mice, in contrast to the expression previously described by splenic microvascular endothelial cells (**Figure 7**). Longitudinal sections from Tamoxifen-injected Ai14^flox/flox^; vWF-iCre/+ SJL F2 mouse spleens showed absent tdT expression in lymphatic endothelium (white arrows) at low magnification (**Figure 10M-P**), further supported by higher magnification images (**Figure 10Q-X**).

**Figure 10.**
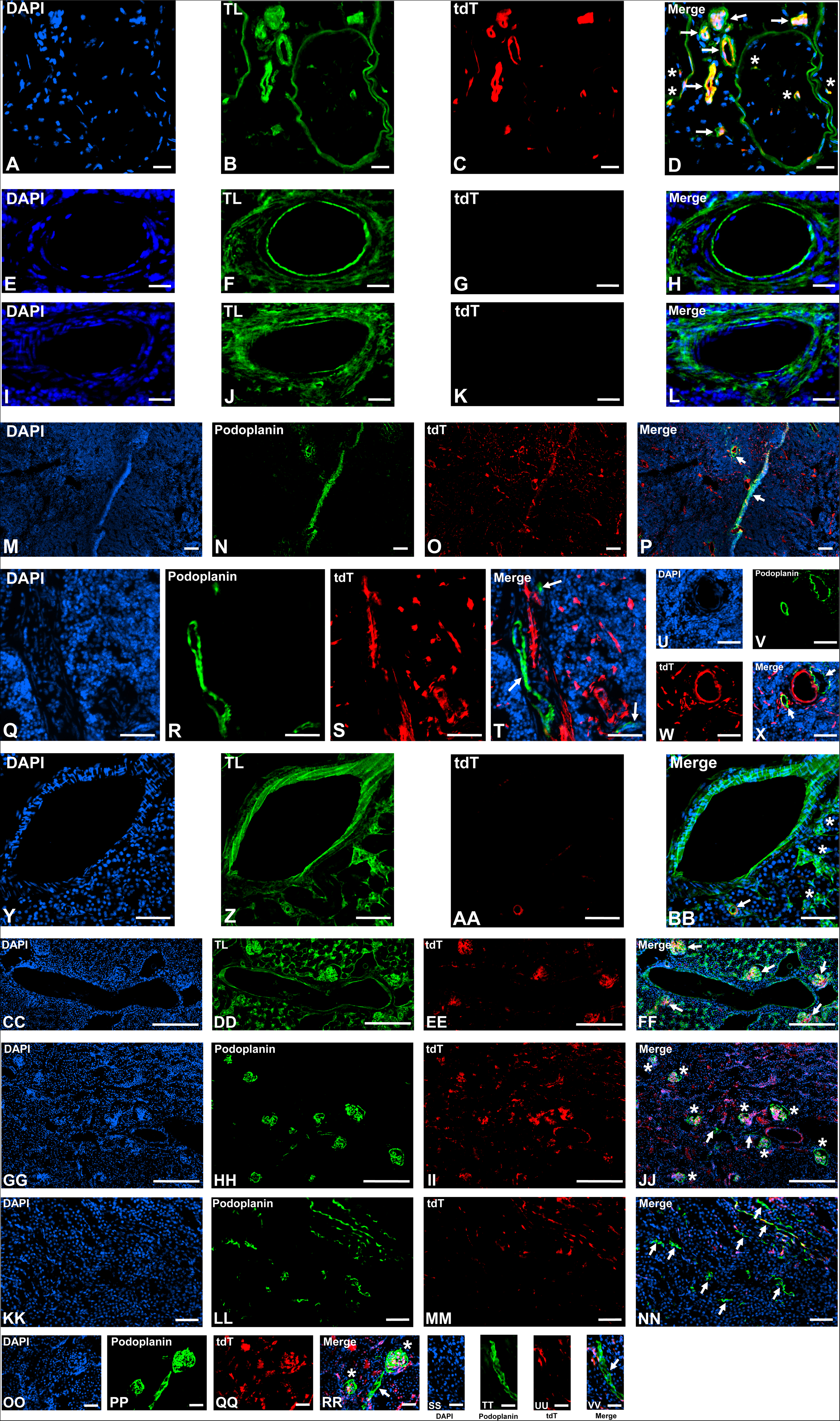
Tamoxifen-induced Cre recombinase expression in large vascular and lymphatic endothelium. Representative digital photomicrographs of paraformaldehyde-fixed, sucrose cryopreserved cryostat axial sciatic nerve sections from an Ai14^flox/flox^; vWF-iCre/+ SJL F2 mouse shows tdT expression co-localizing with TL green fluorescence in epineurial arteriole and venule endothelial cells with diameters < 50 µm (white arrows) in addition to endoneurial microvascular endothelial cells (asterisk) following Tamoxifen-induced recombination (**A-D**). Cre expression is absent from a large splenic artery (**E-H**), vein (**I-L**) and lymphatic vessels (**M-X**, white arrows) on longitudinal sections from Tamoxifen-injected Ai14^flox/flox^; vWF-iCre/+ SJL F2 mice. Similarly, longitudinal sections from Ai14^flox/flox^; vWF-iCre/+ SJL F2 mouse kidneys show absent tdT expression on a TL-reactive large renal cortical artery and peritubular capillaries (asterisk), with expression seen by an adjacent post-capillary venule (white arrow, **Y-BB**) following Tamoxifen-induced recombination. Absent tdT expression is observed in a large TL- reactive renal cortical vein with tdT expression seen on adjacent glomerular capillary endothelial cells (white arrows, **CC-FF**). tdT expression is also absent from renal lymphatic endothelium from the cortex (**GG-JJ**, white arrows) and medulla (**KK-MM**) on longitudinal kidney sections from Tamoxifen-treated Ai14^flox/flox^; vWF-iCre/+ SJL F2 mice, supported by higher magnification images (white arrows**, OO-VV**). Podoplanin-expressing glomerular epithelial cells (green fluorescence) surrounding tdT-expressing capillary endothelial cells are also seen in the cortex at low (**GG-JJ**, asterisk) and higher (**OO-RR**, asterisk) magnification Scale bars: A-L = 50 µm, M-P = 250 µm, Q-BB = 100 µm, CC-FF = 250 µm, GG-RR = 100 µm.

Similarly, longitudinal sections from Tamoxifen-injected Ai14^flox/flox^; vWF-iCre/+ SJL F2 mouse kidneys showed absent or low tdT expression in large renal arteries, in contrast to expression in post-capillary venules, as shown in **Figure 10Y-BB**. As previously shown in **Figure 8**, tdT expression was absent form peritibular capillaries (asterisk), consistent with the known renal microvascular endothelial cell vWF heterogeneity [35]. tdT expression was also absent in large renal veins, in contrast to glomerular capillaries (**Figure 10CC-FF**, white arrows). As described in the spleen, longitudinal sections from Tamoxifen-injected Ai14^flox/flox^; vWF-iCre/+ SJL F2 mouse kidneys showed absent tdT expression from lymphatic endothelium from the cortex (**Figure 10GG-JJ**, white arrows) and medulla (**Figure 10KK-NN**, white arrows), supported by higher magnification images from the cortex (**Figure 10OO-RR**, white arrows) and medulla (**Figure 10SS-VV**) respectively. Podoplanin-expressing glomerular epithelial cells surrounding and partly overlapping tdT-expressing capillary endothelial cells (asterisk) were seen at low and higher magnification as shown in **Figure 10GG-JJ and OO-RR** respectively. These observations from large arteries and veins, and lymphatic vessels from several organs collectively support Tamoxifen-induced Cre recombinase expression driven by a microvascular endothelial-cell selective vWF promoter in Tamoxifen-injected Ai14^flox/flox^; vWF-iCre/+ SJL F2 mice.

Tamoxifen-dependent **(Figure 11A-F**) and -independent tdT expression (data not shown) was observed in the bone marrow megakaryocytes of Ai14^flox/flox^; vWF-iCre/+ F2 SJL mice, with absent tdT expression observed in Ai14^flox/flox^; +/+ F2 SJL control mice megakaryocytes (**Figure 11G-L**). As previously shown in **Figure 7**, reduced tdT expression was observed in splenic megakaryocytes compared to microvascular endothelial cells in Tamoxifen-induced Ai14^flox/flox^; vWF-iCre/+ SJL F2 mice, supporting the notion that the transgene was preferentially transcribed in microvascular endothelial cells. However, variable Tamoxifen-independent microvascular endothelial cell Cre expression was also observed in the organs of some Ai14^flox/flox^; vWF-iCre/+ mice evaluated (highest in the spleen and lowest in the brain [data not shown]), indicative of transient Cre recombinase expression during development or postnatally in corn oil-treated mice or germline transmission of the excised tdT allele to these mice. This has been previously described as a significant limitation of the Ai14^flox/flox^ mouse strain to report Cre expression. This is as a consequence of the short distance between loxP sites flanking the stop codon, which has been shown to reduce recombination efficiency [6, 7, 37]. Despite these limitations, these data show that the vWF-iCre transgene is preferentially expressed in microvascular endothelial cells and megakaryocytes in the SJL background. However, independent recombination events preclude the use of Ai14 mice to report time-dependent microvascular endothelial cell specific gene deletion in vWF-iCre/+ SJL mice.

**Figure 11.**
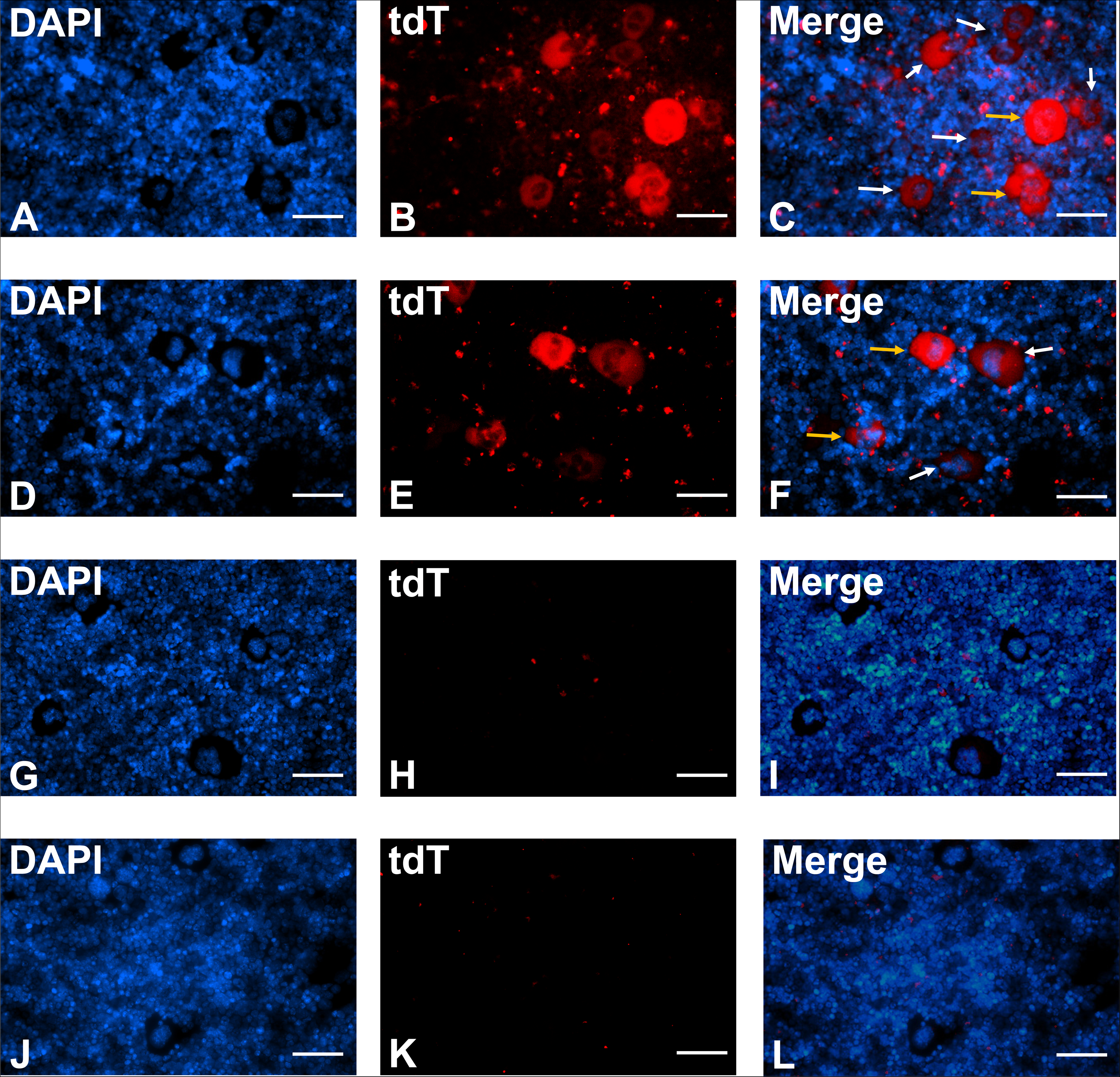
Tamoxifen-induced Cre recombinase expression in bone marrow megakaryocytes of Ai14^flox/flox^; vWF-iCre/+ SJL F2 mice. Representative digital photomicrographs of paraformaldehyde-fixed bone marrow smears from Ai14^flox/flox^; vWF-iCre/+ F2 mice show high (yellow arrows) and low (white arrows) megakaryocyte-specific tdT expression following Tamoxifen-induced recombination (**A-F**) that does not occur in control mice lacking the vWF- iCre transgene (**G-L**). Scale bars: A-L = 50 µm.

## Discussion

We have successfully developed a Tamoxifen-inducible vWF Cre recombinase SJL mouse strain with high recombination efficiency demonstrated in bone marrow megakaryocytes, and specific expression in microvascular endothelial cells in multiple organs, including peripheral nerves, using two reporter mouse strains. A single transgene copy without structural variants is stably integrated into chromosome 1 with deletion of a 125 kb genomic fragment that encodes *Cntn2* (National Library of Medicine National Center for Biotechnology Information [NLM-NCBI] Gene ID: 21367) and *Nfasc* (NLM-NCBI Gene ID: 269116). Cntn2 and Nfasc are members of the immunoglobulin subfamily of cell adhesion molecules required for neuronal axonal guidance during murine cerebellar development. vWF-iCre/+ SJL mice are viable, productive with a normal lifespan and do not show any obvious signs of abnormal behaviors, cerebellar dysfunction or seizures as *Cnt2*^+/-^ and *Nfasc*^+/-^ mice. Furthermore gross and microscopic evaluation of Ai14^flox/flox^; vWF-iCre/+ SJL F2 mouse brains did not show structural differences compared to Ai14^flox/flox^; +/+ F2 SJL control mice (data not shown).

Detailed protocols are provided to allow independent verification of these results, including the vWF-iCre transgene sequence, and the authenticity of this strain has been validated by the University of Missouri MMRRC prior to repository acceptance. SJL mice were chosen for this strain due to our interest in studying the role of time-dependent blood-nerve barrier microvascular endothelial cell gene deletion in the experimental autoimmune neuritis model of Guillain-Barré syndrome. This disorder is more severe and more representative of GBS in SJL mice compared to C57BL/6 mice [38–40]. However, backcross to the more commonly used C57BL/6, or any other mouse background, for experimental purposes is relatively straightforward, using the same primers and PCR protocol described previously for transgene identification.

A *vWF* transgenic FVB mouse strain generated using the similar human microvascular endothelial cell specific construct coupled to an *Escherichia coli LacZ* gene and a simian virus poly(A) sequence was developed ∼30 years ago [20]. Despite demonstrated vWF protein expression on endothelial cells of the spleen, testis, heart, lungs, liver and kidneys by immunohistochemistry, there was a failure to detect β-galactosidase on endothelial cells in those organs [20]. Importantly, the transgene copy number and integration site, as well as transgene construct or LacZ mRNA stability in the generated mouse lines were not reported. Furthermore, potassium ferri- and ferro-cyanide-based X-gal staining methods to detect β- galactosidase are more efficient in detecting expression in embryos compared to thin organ/ tissue sections [41], implying that a failure to detect microvascular endothelial cell β- galactosidase in several organs could also have been a technical limitation. Moreover, this mouse strain is not commercially available to our knowledge, providing us with the impetus to develop and validate an inducible microvascular endothelial cell specific mouse strain based on Cre-loxP methods, and demonstrate specific Cre recombinase expression using commercially available validated mouse reporter mouse strains, for widespread scientific community use.

The observed Tamoxifen-independent megakaryocyte and microvascular endothelial cell Cre expression in some Ai14^flox/flox^; vWF-iCre/+ mice that was maximal in the spleen and least observed in the brain supports the notion that Cre expression is cell-specific, with potential transient Cre expression during development or postnatally accounting for these observations [6, 7]. No unexpected Cre expression occurred in the selected vWF-iCre/+ SJL mouse organs studied using the Ai14 strain apart from gastrocnemius muscles that exhibited a variable mosaic pattern in fascicles in both Tamoxifen- and corn-oil treated mice. Unexpected expression of Cre recombinase may occur due to transient expression of genes in the germline or during early development [6, 7], suggesting possible myogenic vWF gene transcription with transient Cre-mediated recombination.

As discussed previously, no functional murine myogenic bHLH motifs were detected in the vWF- iCre transgene sequence. Importantly, we observed germline recombination of the *H2-Aa* 5’ and 3’ loxP sites, as detected by a three-primer genotyping strategy [6], with our recently developed Major Histocompatibility Compatibility class II A-alpha subunit conditional knockout SJL mouse (H2-Aa^flox/flox^; officially designated as SJL.B6-H2-*^Aatm1c(KOMP)Wtsi^*/UbeeMmmh; RRID:MMRRC_071292-MU) in mixed strain H2-Aa^flox/+;^ Ai14^flox/+^ SJL mice with and without the vWF-iCre transgene. We have not observed any Tamoxifen-independent floxed allele recombination events in >30 generations of H2-Aa^flox/flox^; vWF-iCre/+ or H2-Aa^flox/flox^; +/+ SJL mice to date (data not shown). This implies a potential for some nonspecific endogenous CAG promoter-driven Cre recombinase expression intrinsic to Ai14 mice during development. These observations imply that Ai14 mice are inadequate to study the dynamics of vWF-iCre transgene expression in conditional gene knockout or knock in these SJL mice.

Currently used mouse vascular endothelial cell Cre lines (e.g. Tie2-CreERT2, Cdh5-CreERT2) target either all vascular endothelial cells or are not endothelial cell-restricted [1]. Furthermore, evaluation of peripheral nerve vascular targeting is also lacking from these cell lines. Based on single cell transcriptome data from the mouse Sciatic Nerve Atlas (SNAT: https://www.snat.ethz.ch) [18], several mouse lines that target brain endothelial cells failed to show transcript expression in sciatic nerve endothelial cells (e.g. Sftpa1-CreERT2), are expressed at high levels in multiple peripheral nerve cells (e.g. Acvrl1-CreERT2), at high levels in endothelial cells and low levels in multiple peripheral nerve cells such as Schwann cells and perineurial cells (e.g. Mfsd2a-CreERT2), or at high levels in endothelial cells and the perineurium (Slco1c1-CreERT2). Although, transcript expression does not necessarily imply functional protein translation, vWF protein expression occurs only in microvascular endothelial cells in multiple organs and megakaryocytes in humans and mice [19, 20]. Despite the limitations of the Ai14 mouse, the very low or absent f tdT expression observed in large arteries, veins and lymphatic vessels implies that the vWF-iCre/+ SJL mouse is useful for targeting microvascular endothelial gene alterations in multiple organs such as peripheral nerves, brain, spleen, kidney and muscle. Further studies are needed to verify microvascular endothelial cell specificity with more robust reporter mouse strains, as well as determine induced Cre recombinase expression in other organs such as the skin, eyes, lungs, heart, gut and testis.

Studies evaluating specific gene alterations in microvascular endothelial cells using the vWF- iCre/+ SJL mice should consider whether the gene(s) of interest is/are constitutively expressed in bone marrow or splenic megakaryocytes to ensure that platelet formation and function are not disrupted, as this could compromise wound healing or result in unexpected phenotypes. SJL mice are not great breeders, so these mice may be backcrossed to better breeder strains, such as the commonly used C57BL/6 background based on the experimental model studied. vWF- iCre/+ mice would support ongoing studies aimed to understand the junctional complex and transporter biology of restrictive tight junction-forming microvascular endothelial barriers, such as the blood-brain, blood-nerve, blood-retina and blood-testis barriers during development, in health and normal aging, and in different disease states, guided by observational protein, proteome, transcript or transcriptome data from human specimens. Of particular relevance to peripheral nerves, the vWF-iCre model provides an avenue to study undetermined key molecular determinants and signaling pathways of unique biologic networks ascertained from the published human blood-nerve barrier transcriptome, including microvascular endothelial cell-to-cell, and endothelial cell-glial, -pericyte, -perineurial and -immune cell crosstalk, to mention a few [42].

## Conclusions

A viable Tamoxifen-inducible vWF Cre recombinase SJL mouse strain has been developed with stable integration of a single transgene copy without structural variants in chromosome 1 and microvascular endothelial cell-specific expression in multiple organs, including peripheral nerves. These mice may undergo backcrossing to different mouse backgrounds without transgene alteration. This Cre recombinase mouse strain would support hypothesis-driven mechanistic studies to decipher the role(s) of specific genes transcribed by microvascular endothelial cells during development, as well as in physiologic and pathophysiologic states in an organ- and time-dependent manner. Careful selection of reporter mouse strains that do not exhibit significant Tamoxifen-independent Cre expression or independent random recombination events around floxed alleles is essential to track the dynamics of conditional gene deletion or expression *in vivo* or *in situ* using this strain. Work is ongoing to decipher the essential or redundant role(s) of specific junctional complex genes transcribed by the restrictive tight-junction forming blood-nerve barrier endoneurial endothelial cells during development and following traumatic nerve injury using this model.

## Acknowledgments

We acknowledge Wanda Filipiak and Galina Gavrilina for generating founder transgenic mice via pronuclear microinjection of fertilized eggs and the Transgenic Animal Model Core of the University of Michigan’s Biomedical Research Core Facilities for their technical expertise. Special thanks to past members of the Neuromuscular Immunopathology Research Laboratory, particularly Dr. Chaoling Dong, for their essential technical assistance utilized to develop, characterize, expand and maintain the Tamoxifen-inducible von Willebrand factor Cre recombinase SJL mouse strain colony at the University of Alabama at Birmingham. We acknowledge the University of Missouri Mutant Mouse Research & Resource Center (MMRRC) and Cergentis B.V. for performing transgene copy number and chromosomal integration site and integrity analyses on our inducible von Willebrand factor Cre recombinase SJL mouse strain respectively.

## Data Statement

Data that support the study’s findings are available to members of the scientific community on request from the corresponding author, E.E.U. Data pertaining to this mouse strain are also publically achieved at https://doi.org/10.1101/2023.07.24.550419 and https://doi.org/10.1101/2023.07.24.550421.

**Supplementary File 1.** vWF-iCre transgene sequence. The confirmed transgene sequence containing the human vWF promoter, with the microvascular endothelial cell-selective sequence located at position 236 to 969 (yellow highlight), followed by the iCre sequence and the bgh-polyA signal is shown.

**Supplementary Figure 1.** Mixed strain C57BL/6-SJL vWF-iCre/+ founder and reporter mice genotyping. Reverse digital photomicrographs of 2% agarose electrophoresis gels in 1X TAE buffer show tail snip genotyping results following crossing with mT/mG^flox/flox^ mice. mT/mG^flox/+^; vWF-iCre/+ F1 mice were generated to further determine Cre recombinase expression cell specificity and efficiency. The vWF-iCre transgene is identified at 604 bp, with the mutant and wildtype *mT/mG* identified at 128 bp and 212 bp respectively (**A**). Representative reverse digital photomicrographs of 2% agarose electrophoresis gels in 1X TAE buffer tail snip genotyping results following congenic SJL vWF-iCre/+ mouse crossbreeding with Ai14^flox/flox^ mice, followed by mixed strain Ai14^flox/+^; vWF-iCre/+ F1 mice bred with Ai14^flox/flox^ mice to generate Ai14^flox/flox^; vWF-iCre/+ F2 mice to further determine Cre recombinase cell expression specificity. The vWF-iCre transgene is identified at 604 bp, with the mutant and wildtype *Ai14* identified at 196 bp and 297 bp respectively (**B**). C represents lanes with absent DNA (ddH_2_O) template controls.

**Supplementary Figure 2.** Comparative Tamoxifen-induced Cre recombinase expression on sciatic nerve endoneurial microvessels in Ai14^flox/flox^; vWF-iCre/+ SJL F2 mice. Representative digital photomicrographs of paraformaldehyde-fixed, sucrose cryopreserved sciatic nerve axial sections from the same mouse show similar tdT co-localization on vascular endothelial cells detected by TL (**A-D**, white arrows) and CD31 (**E-H**, white arrows), with more intense perineurial staining seen with TL compared to CD31 (asterisk in **B**).

