## Supplemental File 1 for "A novel inducible von Willebrand Factor Cre Recombinase mouse strain to study microvascular endothelial cell-specific biological processes *in vivo*"

5' GGCCGCGTTTaaacATTAGGCTTTTGGGAATGGGAGGCTGGGAAGAAGAATTGTGTGTGT 60

PmeI | VWF human promoter
notI | PmeI

5' GTGTGTGTGTGTGTGTGTGTGTGTGTGTGTGTGTGTAAGATCAGGGTACCAGAAGTGGGTGGAA 120

VWF human promoter

5' ATGTCCTTGAGAATTAGAATTATTAGAATGTAGCAACAGTAGAAGTATTAGACTCAAACC 180

VWF human promoter

5' ATCACTCCCCACCTTCACCATTTTACAAAGGCTTAGGCTTGTGGCCAAGACCTTCATCTT 240

VWF human promoter

5' TAGCCGATCCATTCAACCCTGGCCAGGATCCAAATGGACTGTTTTTGTCAGGGCCAGGAC 300

VWF human promoter

5' CGGATCCTTCATACCTGGGGTGCATAGGAAGTGTTAGTACTCCCCTTCCTCCAAACACAG 360

VWF human promoter

5' CAGCAAAATTGGCTCAGGTTGAGGTGTTTTTCTCAACTTCCCTGGAGTCCAGCCCTGGAA 420

VWF human promoter

5' GCTGGATCAGGAAGCTGTGTTGTTCTACTGTGATTCCCCCTGGCCTGTATCAGCTTGCCC 480

VWF human promoter

5' TGAAACAACCAGCATTCCTGGTTATCCCACACAGGTGGGGCACTCTAGGAAGACCAGGGA 540

VWF human promoter

5' TCAAGTGTGGGGGTGTAGGGATAGGGGGTGTTTGGGGAGGGCAAGGCAGTTAATTAAGGC 600
VWF human promoter

5' AGCTGCCAGGAGGTCTCCCTCCAAACTCTACAAAGCTTTATCAGCTTGGAGGTACTTCTA 660
VWF human promoter

5' ATACCATTTCCTTTCATTGTTTCCTTTTGGTAATTAAAAGGAGGCCAATCCCCTGTTGTG 720
VWF human promoter

5' GCAGCTCACAGCTATTGTGGTGGAAAGGGAGGGTGGTTGGTGGATGTCACAGCTTGGGC 780
VWF human promoter

5' TTTATCTCCCCCAGCAGTGGGGACTCCACAGCCCCTGGGCTACATAACAGCAAGACAGTC 840
VWF human promoter

5' CGGAGCTGTAGCAGACCTGATTGAGCCTTTGCAGCAGCTGAGAGCATGGCCTAGGGTGGG 900
VWF human promoter

5' CGGCACCATTGTCCAGCAGCTGAGTTTCCCAGGGACCTTGGAGATAGCCGCAGCCCTCAT 960
VWF human promoter

5' TTGCAGGGGAAGGTATGGCCTTTGGAAGGAGAGCTGGCTCAGTTGTGGGAGGAAGATGCA 1020
VWF human promoter

5' GGACTGACTGATCCCTGCTCCTGGGGAGCTGGAGTTCTCTGTCGCTGGACTAGAAGGGCT 1080
VWF human promoter

5' TTGTTTGGAGGGGCAATTCAATTCAGCCAGGGATGATCCTAATACTCCCTCCTCCACTTG 1140

VWF human promoter

5' CCTCTGAGGGTCCTGGGGCTGCTTTTCTTCATGCAGTGGGTTTTACTGTTTGATAGTACT 1200

VWF human promoter

5' TCACTCAAATGAGTTGGAATGAAGTTTGCCCTCACCTCTGAGAACCTGGGAGCAGCTGAA 1260

VWF human promoter

5' TGTACCTGCGTGTTAGGACTGGGAGGGGACACCTGCTTGGAGACCGAGACCTGGCAGTAT 1320

VWF human promoter

5' CTGACATCTCAGTGTTCCTTCCACAGATGTATCACAGATTGGCTTGATTTCACCTTTGGC 1380

VWF human promoter

5' TGGATGGGACCTTAGGTAGGAAGGGAGTCACCCCCAGTGAATCTCAGGCAGCAGATTCTG 1440

VWF human promoter

5' CACTTCATTTAACAACTTTTCCCGAGGAGAGGGGCTACAGCAGGGGCTCTAAGTGACTTG 1500

VWF human promoter

5' GGGTACGCTCTGCCAGCCAGGATGAATTGTCCCTCTCTTGGGGGTCACACAGTGGGGAAG 1560

VWF human promoter

5' TCTGCCTGCATCCAGGGCCGCTGGACTCCTGTCCATTTTTTCAGATGAACTCAGCAAACA 1620

VWF human promoter

5' TTTGCTGGGCATCTCCTGGGTGCTAAGCATCTTGCCAGGTGCTGGGGTTGGAGGCAAGGG 1680

VWF human promoter

5' AGACAGCCTTTGCTCTTGTGAAGGCACTTGTGGTACAGAGTCAGGGGCCAACAAGCAAAC 1740

VWF human promoter

5' CGTCAAGTTGGTGGTTCCTGAGCATTCTCTATGTCTGGGCTGCTGTGGTGGGCACACAAG 1800

VWF human promoter

5' TGTAAGACGGTTCCTACTCGCCAGTTTGGATGCAGAGGCAGGAAGGAATGAGGTGTGTGT 1860

VWF human promoter

5' TAGCTCCCAGCTGCTTCAGGAGGCAGGGATGTGAGGCCCAGCGGGCCTGGAGGGAAGGCA 1920

VWF human promoter

5' GCGTTTTCCTCCTGTCTTGGGCCTGGGACTGCTGTCTGTGGAAAGGTGCCCACAGGTCCC 1980

VWF human promoter

5' AGCTCACAGCGATTGTTACCCTTGGGCCTGGCACTGGCCAGGGGTTTTTCGGGGGCCAG 2040

VWF human promoter

5' AAGTCCATGTTCAAAGGGGAAAAGGGGGTCACGAGGATCAATCTTTTCTCCTGCTTTAAA 2100

VWF human promoter

5' GAAATGTTTTTGCTACTGCATGCCCTGATAGTCGCCACACCAGCAGCCGCCTACCTGGGC 2160

VWF human promoter

5' AGCAATGACCAGCTCACGTCTCTTGCTTCTTTGCAGATGGTGCCCAAGAAGAAGAGGAAA 2220

VWF human promoter | iCre

M V P K K K R K
1 2 3 4 5 6 7 8

5' GTCTCCAACCTGCTGACTGTGCACCAAAACCTGCCTGCCCTCCCTGTGGATGCCACCTCT 2280

iCre

ApaLI

V S N L L T V H Q N L P A L P V D A T S
9 10 11 12 13 14 15 16 17 18 19 20 21 22 23 24 25 26 27 28

5' GATGAAGTCAGGAAGAACCTGATGGACATGTTCAGGGACAGGCAGGCCTTCTCTGAACAC 2340

iCre

D E V R K N L M D M F R D R Q A F S E H
29 30 31 32 33 34 35 36 37 38 39 40 41 42 43 44 45 46 47 48

5' ACCTGGAAGATGCTCCTGTCTGTGTGCAGATCCTGGGCTGCCTGGTGCAAGCTGAACAAC 2400

iCre

T W K M L L S V C R S W A A W C K L N N
49 50 51 52 53 54 55 56 57 58 59 60 61 62 63 64 65 66 67 68

5' AGGAAATGGTTCCCTGCTGAACCTGAGGATGTGAGGGACTACCTCCTGTACCTGCAAGCC 2460

iCre

R K W F P A E P E D V R D Y L L Y L Q A
69 70 71 72 73 74 75 76 77 78 79 80 81 82 83 84 85 86 87 88

5' AGAGGCCTGGCTGTGAAGACCATCCAACAGCACCTGGGCCAGCTCAACATGCTGCACAGG 2520

iCre

R G L A V K T I Q Q H L G Q L N M L H R
89 90 91 92 93 94 95 96 97 98 99 100 101 102 103 104 105 106 107 108

5' AGATCTGGCCTGCCTCGCCCTTCTGACTCCAATGCTGTGTCCCTGGTGATGAGGAGAATC 2580

iCre

R S G L P R P S D S N A V S L V M R R I

109 110 111 112 113 114 115 116 117 118 119 120 121 122 123 124 125 126 127 128

5' AGAAAGGAGAATGTGGATGCTGGGGAGAGAGCCAAGCAGGCCCTGGCCTTTGAACGCACT 2640

iCre

R K E N V D A G E R A K Q A L A F E R T

129 130 131 132 133 134 135 136 137 138 139 140 141 142 143 144 145 146 147 148

5' GACTTTGACCAAGTCAGATCCCTGATGGAGAACTCTGACAGATGCCAGGACATCAGGAAC 2700

iCre

D F D Q V R S L M E N S D R C Q D I R N

149 150 151 152 153 154 155 156 157 158 159 160 161 162 163 164 165 166 167 168

5' CTGGCCTTCCTGGGCATTGCCTACAACACCCTGCTGCGCATTGCCGAAATTGCCAGAATC 2760

iCre

L A F L G I A Y N T L L R I A E I A R I

169 170 171 172 173 174 175 176 177 178 179 180 181 182 183 184 185 186 187 188

5' AGAGTGAAGGACATCTCCCGCACCGATGGTGGGAGAATGCTGATCCACATTGGCAGGACC 2820

iCre

R V K D I S R T D G G R M L I H I G R T

189 190 191 192 193 194 195 196 197 198 199 200 201 202 203 204 205 206 207 208

5' AAGACCCTGGTGTCCACAGCTGGTGTGGAGAAGGCCCTGTCCCTGGGGGTTACCAAGCTG 2880

iCre

K T L V S T A G V E K A L S L G V T K L

209 210 211 212 213 214 215 216 217 218 219 220 221 222 223 224 225 226 227 228

5' GTGGAGAGATGGATCTCTGTGTCTGGTGTGGCTGATGACCCCAACAACTACCTGTTCTGC 2940

iCre

V E R W I S V S G V A D D P N N Y L F C
229 230 231 232 233 234 235 236 237 238 239 240 241 242 243 244 245 246 247 248

5' CGGGTCAGAAAGAATGGTGTGGCTGCCCCTTCTGCCACCTCCCAACTGTCCACCCGGGCC 3000

iCre

R V R K N G V A A P S A T S Q L S T R A
249 250 251 252 253 254 255 256 257 258 259 260 261 262 263 264 265 266 267 268

5' CTGGAAGGGATCTTTGAGGCCACCCACCGCCTGATCTATGGTGCCAAGGATGACTCTGGG 3060

iCre

L E G I F E A T H R L I Y G A K D D S G
269 270 271 272 273 274 275 276 277 278 279 280 281 282 283 284 285 286 287 288

5' CAGAGATACCTGGCCTGGTCTGGCCACTCTGCCAGAGTGGGTGCTGCCAGGGACATGGCC 3120

iCre

Q R Y L A W S G H S A R V G A A R D M A
289 290 291 292 293 294 295 296 297 298 299 300 301 302 303 304 305 306 307 308

5' AGGGCTGGTGTGTCCATCCCTGAAATCATGCAGGCTGGTGGCTGGACCAATGTGAACATa 3180

iCre

R A G V S I P E I M Q A G G W T N V N I
309 310 311 312 313 314 315 316 317 318 319 320 321 322 323 324 325 326 327 328

5' GTGATGAACTACATCAGAAACCTGGACTCTGAGACTGGGGCCATGGTGAGGCTGCTCGAG 3240

iCre

V M N Y I R N L D S E T G A M V R L L E
329 330 331 332 333 334 335 336 337 338 339 340 341 342 343 344 345 346 347 348

5' GATGGGGACTGAACGCGTAGCGGCCCTAGAGCTCGCTGATCAGCCTCGACTGTGCCTTCT 3300

iCre
bgh-pA
BclI
MluI

D G D .
349 350 351 352

5' AGTTGCCAGCCATCTGTTGTTTGCCCCTCCCCCGTGCCTTCCTTGACCCTGGAAGGTGCC 3360

bgh-pA

5' ACTCCCACTGTCCTTTCCTAATAAAATGAGGAAATTGCATCGCATTGTCTGAGTAGGTGT 3420

bgh-pA

5' CATTCTATTCTGGGGGGTGGGGTGGGGCAGGACAGCAAGGGGGAGGATTGGGAAGACAAT 3480

bgh-pA

5' AGCAGGCATGCTGGGGATGCGGTGGGCTCTATGGCTTCTGAGGCGGAAAGAACCACATAT 3540

bgh-pA
NdeI

5' GGC 3543
