## Supplementary figures and images for "A novel inducible von Willebrand Factor Cre Recombinase mouse strain to study microvascular endothelial cell-specific biological processes *in vivo*"

### Supplemental Figure 1

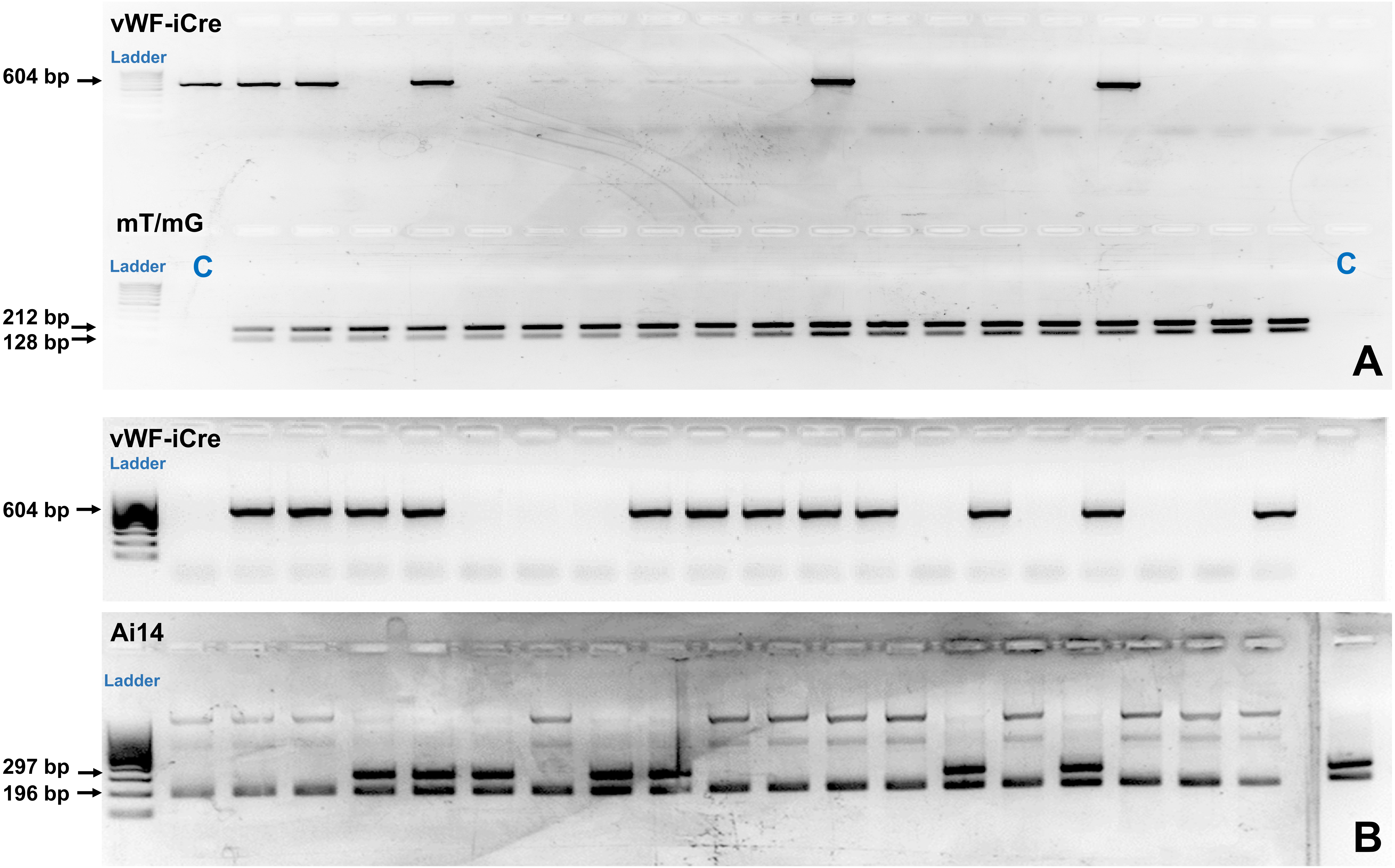

### Supplemental Figure 2

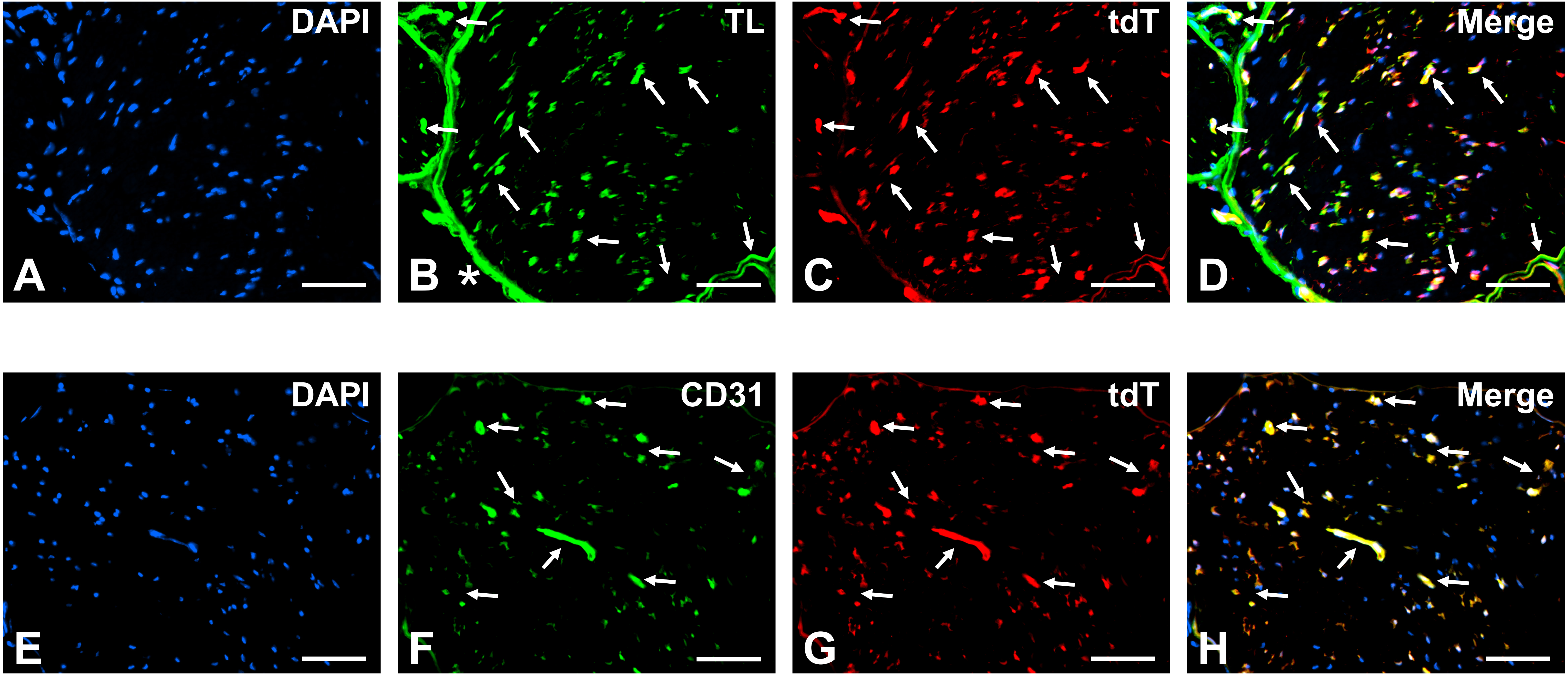
